# ALS sensitive spinal motor neurons enter a degenerative downward spiral of impaired splicing and proteostasis

**DOI:** 10.1101/2022.03.26.485939

**Authors:** Dylan E. Iannitelli, Albert Tan, Emma Nguyen, Asha Babu, Setiembre Delfina Elorza, Tyler Joseph, Sophie Zaaijer, Disi An, Michael Ward, Esteban O. Mazzoni

## Abstract

Despite clear therapeutic potential, the mechanisms that confer differential neuronal sensitivity are not well understood. During Amyotrophic Lateral Sclerosis (ALS), sensitive spinal motor neurons (SpMN) die while a subset of rostral cranial motor neurons (CrMN) survive. In this work, we optimized a protocol to differentiate CrMNs and SpMNs from human induced pluripotent stem cells (iPSCs) by direct programming and positional patterning. Human iCrMNs are more resistant than iSpMNs to proteotoxic stress and rely on the proteasome to maintain proteostasis. iCrMNs better prevent mislocalization of TDP43 from the nucleus, a hallmark of ALS progression. iSpMNs contain more splicing defects than iCrMNs in response to ALS-related stress with genes involved in splicing and proteostasis maintenance. Therefore, iCrMNs resist ALS at two levels, preventing protein accumulation and reducing splicing defects in response to TDP43 nuclear depletion. Thus, ALS-sensitive iSpMNs appear to enter a downward spiral compromising their ability to maintain proteostasis and splicing.

## Introduction

Neurodegenerative diseases are characterized by the selective death of specific populations of neurons. A fundamental question in neurodegenerative disease pathology is why some neurons are susceptible to disease progression and die while other neurons are resistant and survive. Despite the clear therapeutic potential, there are few studies aimed toward the molecular mechanisms underlying differential sensitivity to neurodegeneration. In Amyotrophic Lateral Sclerosis (ALS), most spinal motor neurons (SpMN) progressively degenerate leading to muscle denervation and eventual paralysis and death^1^. However, a subset of cranial motor neurons (CrMN, oculomotor, trochlear, and abducens motor neurons) controlling eye movements typically survive and remain functional in patients until late disease stages^2–4^. Key insights into ALS disease pathology and its potential therapies lay in understanding the differential sensitivity between CrMNs and SpMNs.

The selective resistance of CrMNs compared to SpMNs has been found in a mouse model of ALS expressing human SOD1 G93A and in mouse ESC derived *in vitro* models of oculomotor/CrMN and SpMN differential sensitivity to ALS^5–7^. Our recent work uncovered a superior ability of mouse ESC-derived CrMNs to maintain proteostasis compared to SpMNs, possibly owing to higher protein expression of catalytic proteasome subunits and higher proteasome activity^7^.

A key hallmark of ALS pathology in patients is the accumulation of insoluble protein aggregates, with proteostasis impairment and endoplasmic reticulum (ER) stress factors in neuronal sensitivity to ALS and other neurodegenerative diseases^8–11^. Nearly 90% of ALS cases are understood to be sporadic while familial mutations account for the remaining 10%. ALS causing mutations often lead to misfolding and aggregation and subsequent loss of function of the affected protein and/or toxic gain of function of the resulting misfolded protein. Such is the case for SOD1, FUS, C9ORF72, and TDP43^12–15^. TDP43 plays a functional role in splicing, transcriptional regulation, mRNA stability and transport, stress granule formation and microRNA processing^16,17^. Across familial as well as sporadic ALS cases, TDP43 aggregation is typically preceded by cytoplasmic mislocalization out of the nucleus. Nuclear depletion of TDP43 leads to disrupted splicing, with cryptic exon inclusion shown to cause premature polyadenylation and stop-codon truncation in two neuronal synaptic genes, *STMN2* and *UNC13A*^18–23^. However, it remains unclear how nuclear depletion of TDP43 and other RNA binding proteins (RBP) and splicing defects interact with proteostasis impairment and ER stress. Furthermore, the extent to which nuclear depletion of RBPs and altered splicing are present in neurons resistant to ALS progression is unknown.

The development of protocols to differentiate motor neuron fate from human ESCs or induced pluripotent stem cells (iPSCs) has enabled the possibility for clinic-relevant modeling of ALS in a dish^24,25^. However, protocols relying on directed differentiation strategies to recapitulate developmental processes are often long. They required the formation and subsequent dissociation of embryoid bodies producing motor neuron fates combined with other related neuronal and non-neuronal cells, limiting the study of motor neuron intrinsic biology. Direct programming strategies employing inducible transcription factor (TF) expression from stable, integrated cell lines enable consistent, reproducible differentiation of cell fates with increased efficiency^26,27^. Neurogenin2 (Ngn2), Islet1 (Isl1), and Lim-homeobox 3 (Lhx3) (together referred to as NIL) forced expression is sufficient to differentiate SpMN fate while Ngn2, Isl1, and Paired Like Homeobox 2A (Phox2a) (together referred to as NIP) is sufficient to generate CrMN fate^28^. The NIP and NIL TF cassettes in mouse are each single copy insertions engineered into the doxycycline inducible cassette exchange locus^29^. Single copy integration allows for sufficient expression to rapidly (after 2 days) differentiate mouse CrMN or SpMN fate in near purity^28^. To achieve CrMN and SpMN differentiation by direct programming in human iPSCs, De Santis, et al. took advantage of the piggyBAC tranposase system to integrate multiple copies of inducible NIP or NIL into the genome^30^. The differentiation protocol achieves an impressive 90% β3-tubulin positive neuronal culture with NIP induction yielding up to 95% Phox2b positive CrMNs among the pool of β3-tubulin positive neurons.

In this work, we optimized a method to rapidly differentiate rapidly differentiate and positionally pattern human NIP and NIL inducible (i) iCrMNs and iSpMNs in a monolayer culture that allows for high-throughput scalability across plating formats. The optimized protocol differentiated MN fate with over 90% efficiency and specified CrMN or SpMN fate to yield ALS resistant or sensitive cell types. iCrMNs and iSpMNs modeled the differential sensitivity to ALS-related proteotoxic stress and displayed differential cellular phenotypes associated with ALS progression, such as TDP43 aggregation and mislocalization. We also revealed iCrMN-specific reliance on proteasome function to resist ALS-like stress, supporting our previous finding in mouse ESC-derived MNs^7^. For the first time, we found that iCrMNs and iSpMNs differ in the number and the sets of genes with altered splicing after MG132 induced TDP43 nuclear depletion and in response to ALS-inducing TDP43 mutation. Further, iSpMNs displayed specific differential disrupted splicing in genes involved in ubiquitination, proteasome function, and autophagy critical for maintaining proteostasis. These results suggest that iCrMNs resist TDP43 mislocalization and are less affected than iSpMNs to the same levels of TDP43 mislocalization. Therefore, iSpMNs enter a downward spiral dynamic in which proteotoxic stress leads to altered splicing which further exacerbates their ability to maintain proteostasis and unaltered splicing.

## Results

### Rapid, efficient, and scalable direct programming and patterning of Cranial and Spinal Motor Neurons from iPSCs

Consistency in differentiated cell fate for reproducibility across experiments and conditions is essential to generate *in vitro* models of tissues and specific cell types for disease modeling. This consistency and reproducibility in cell fate generation can be achieved through direct programming by inducible TF expression from stable, integrated cell lines^26,27^. However, single-copy integration may not yield sufficient TF expression for efficient cell fate conversion, and viral integration strategies tend to become silenced during differentiation, requiring continued selection in some models^31^. Thus, we opted for a transposon-mediated integration of inducible TF combinations yielding multiple copy insertion followed by selection and rapid screening of differentiating clones (**Figure 1a**). We took advantage of the previously constructed inducible Ngn2-Isl1-Phox2a (NIP) or Ngn2-Isl1-Lhx3 (NIL) piggyBac vectors and selected for stably transposed cells^28,30^. The NIP and NIL plasmids carry Blastidicin resistance, allowing for rapid selection after nucleofection over 5 days. As integration is a unique event in each cell, the pooled population after selection is highly variable with respect to the number and location of stable integrations and TF expression (**Supplemental Figure 1a-b**). We thus picked individual clones to compare against one another and the pooled population after selection. After 2 days of NIP or NIL induction with doxycycline, we measured induction rates by staining with an antibody for Isl1 (**Supplemental Figure 1a**). Clonal optimization yields a dramatic, over two-fold increase in induction efficiency compared to the pooled culture after antibiotic selection (**Supplemental Figure 1b**). We then took clones with high induction rates and differentiated them for 10 days to generate NIP induced iCrMNs and NIL induced iSpMNs. Clones with the highest expression rate of motor neuron marker gene VAChT were expanded (**Figure 1a**). We initially carried out the above procedure using an iPSC line with a VAChT-Cre/tdTomato reporter (VAChT-tdT) for rapid and convenient assessment of efficient differentiation to motor neuron fate when optimizing the protocol^32^. We selected one clone each for NIP and NIL, respectively, which differentiated to motor neuron fate with similarly high efficiency to carry forward.

**Figure 1.**
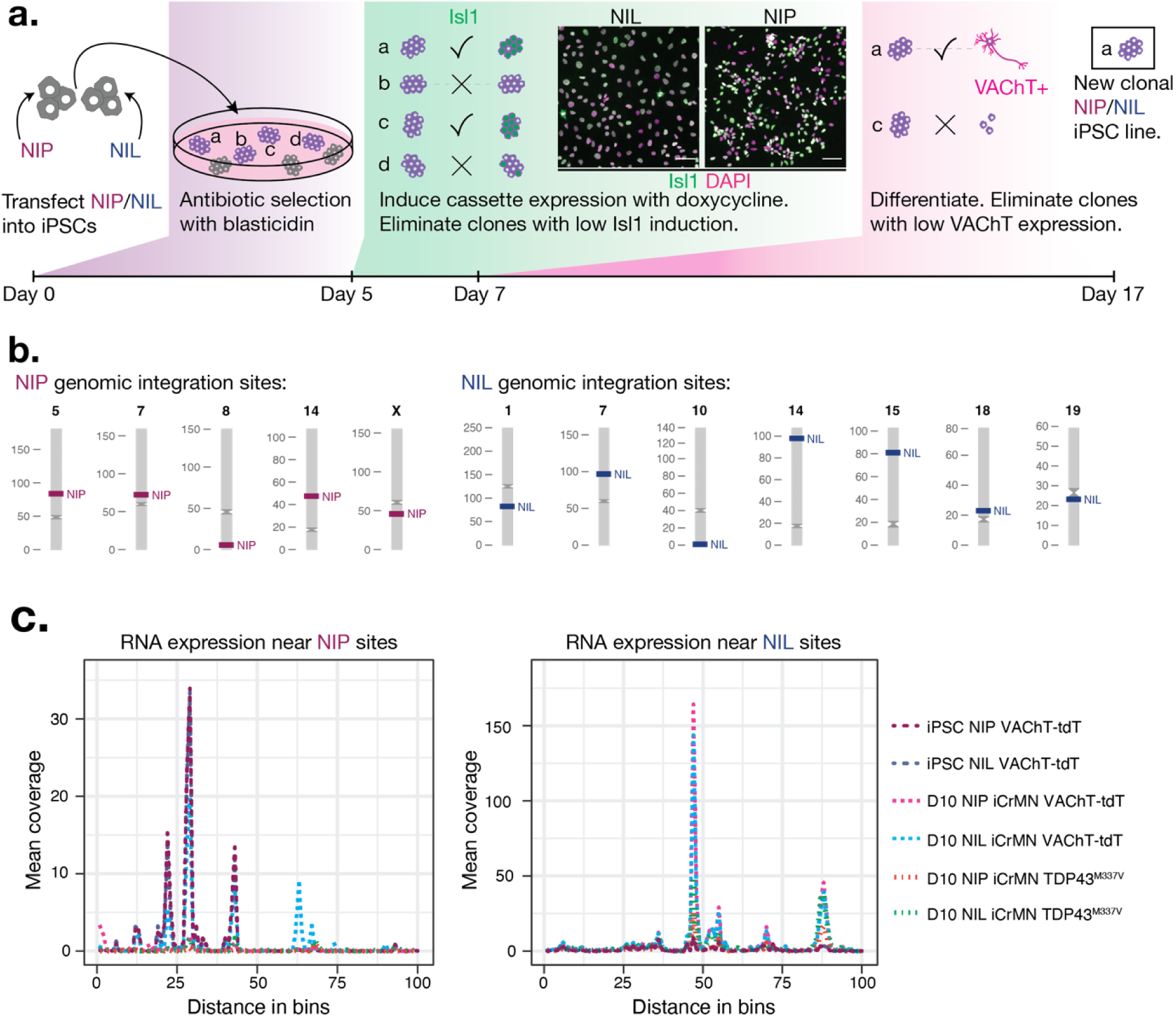
Generating inducible NIP and NIL iPSC lines. (**a**) Schematic for line generation, selection, and screening. iPSCs are transfected by pBAC transposase with NIP or NIL expression cassettes on day 0. From day 0 to day 5, blasticidin selection removes cells without a NIP or NIL insertion. Clones are picked and expanded and NIP and NIL cassette expression is induced for 2 days with doxycycline. Clones with low Isl1 induced expression are eliminated and the rest are differentiated for 10 days. Clones with low VAChT expression are eliminated and those with the highest expression kept as new NIP or NIL inducible iPSC lines. (**b**) Chromosome map of NIP (left, red) and NIL (right, blue) integration sites. (**c**) Metagenome plots. RNA expression near 5 NIP overlayed insertion sites (left) and 7 overlayed NIL insertion sites (right). Windows around insertions and thus bin sizes are variable between insertion sites.

Transposase-based integration strategies result in random integration of the expression vector into the genome. Thus, we performed shotgun whole genome sequencing in collaboration with FindGenomics to identify each NIP or NIL integration site (**Figure 1b**). The NIP line contained 5 and the NIL line 7 independent integration sites spread across the genome. To determine if the expression of nearby genes is affected by the integration of the NIP or NIL cassettes, we looked at nearby mRNA expression in stem cells and day 10 MNs in NIP and NIL VACHT-tdT iPSCs and an independent TDP43 mutant background (**Figure 1c**, **Supplemental Figure 2**). In large part, the presence of an integrated NIP or NIL cassette does not affect nearby transcription, and most integration sites land between genes or inside introns (**Supplemental Figure 2**). We thus established a methodology for rapid generation of inducible motor neuron programming of two motor neuron types with differential sensitivity to neurodegeneration from human iPSC lines.

Traditionally, motor neuron differentiation protocols rely on embryoid body formation after neural induction, making them work-intensive and necessitating cell dissociation before plating neurons^24,25^. We developed and optimized a monolayer differentiation protocol to achieve rapid, highly efficient, and scalable motor neuron cultures across various plating formats (**Figure 2a**). We dissociated cultured NIP or NIL iPSCs and plated them directly into a Neurobasal-N2, B27 based media with doxycycline to induce cassette expression with ROCK inhibitor to improve viability after plating. Two days after plating and induction, previously experimentally determined low (0.01 µM) and high (1.0 µM) concentrations of retinoic acid (RA) positionally pattern rostral CrMN fate and caudal SpMN fate, respectively^33^. 5-Fluoro-20-deoxyuridine (FDU) and Uridine on day 6 eliminate actively dividing cells in the culture. Finally, Vitamin C and neurotrophic factors (BDNF, GDNF, and CNTF) support the viability and long-term culture of motor neurons.

**Figure 2.**
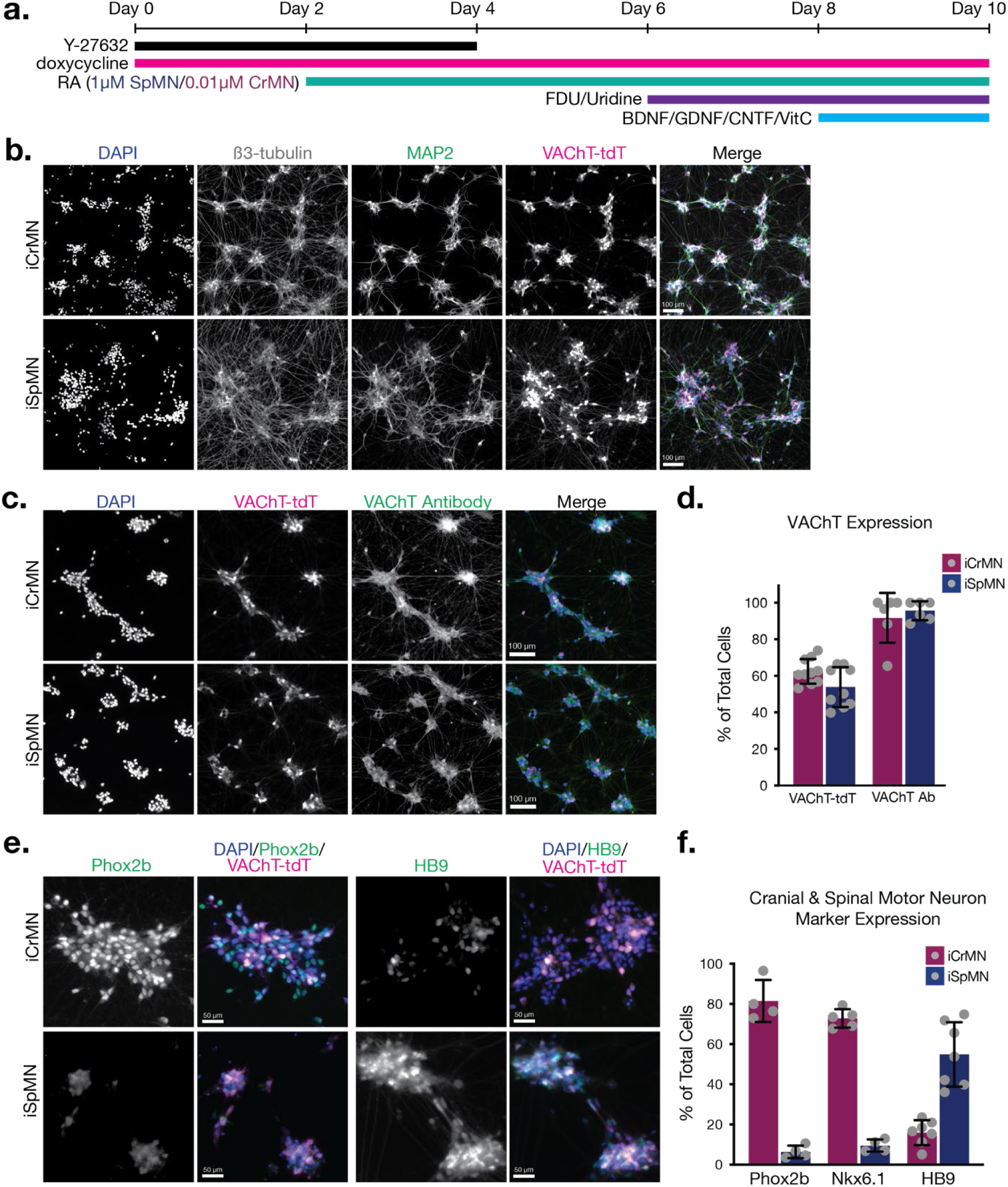
NIP and NIL direct programming and RA patterning differentiate CrMN and SpMN fate. (**a**) Schematic for differentiation protocol. (**b**) Representative images of β3-tubulin, MAP2, and VAChT-tdTomato in day 10 NIP iCrMNs and NIL iSpMNs. Merged image shows DAPI in blue, β3-tubulin in grey, MAP2 in green, and VAChT-tdT in magenta. Scale bar = 100 µm. (**c**) Representative images of VAChT-tdT expression and VAChT antibody staining. Merged image shows DAPI in blue, VAChT-tdTomato in magenta, and VAChT antibody in green. Scale bar = 100 µm. (**d**) Quantification of VAChT-tdTomato expression (n = 10, mean ± SEM) and VAChT antibody (Ab) staining (n = 6, mean ± SEM). (**e**) Representative images of Phox2b and HB9 expression. Merged images show DAPI in blue, Phox2b (second from left) or HB9 (first from right) in green, and VAChT-tdT in magenta. Scale bars = 50 µm. (**f**) Quantification of Phox2b (n = 4, mean ± SEM), Nkx6.1 (n = 5, mean ± SEM), and HB9 (n = 7, mean ± SEM).

As expected, since NIP and NIL contain the pro-neuronal Neurog2 TF, both lines expressed β3-tubulin and MAP2 after 10 days of differentiation (**Figure 2b**). We then asked what proportion of cells become motor neurons by staining for VAChT and CHAT and comparing to the expression of the VAChT- tdTomato reporter (**Figure 2c-d**). While the VAChT- tdTomato reporter was present in 50-60% of both iCrMNs and iSpMNs on average (iCrMN 63.48% ± 6.76, iSpMN 53.77% ± 10.94), 90% or more cells in each culture were positive for the VAChT motor neuron marker (iCrMN 91.74% ± 13.63, iSpMN 95.53% ± 5.16). This discrepancy may be due to stochastic differences in the efficiency of the VAChT- Cre/tdTomato reporter system. iCrMNs express the rostral cranial motor neuron marker genes Phox2b and Nkx6.1 while very few iSpMNs express either (Phox2b: iCrMN 81.46% ± 10.45, iSpMN 6.39% ± 3.10; Nkx6.1: 72.84% ± 4.62, iSpMN 9.54% ± 3.06) (**Figure 2e-f**). On the other hand, around 50% of iSpMNs are positive for HB9 staining with low expression in iCrMNs (see RNA-seq results below; iCrMN 15.94% ± 6.23, iSpMN 52.25% ± 16.64). Together, the high expression of neuronal and motor neuron fate markers indicates that this protocol based on inducible TF expression and patterning signals produces iCrMN and iSpMN within 10 days without replating or dissociation.

### NIP- and NIL-programmed and patterned iPSCs faithfully generate cranial and spinal motor neuron fate, respectively

To deeply assess the specific fates generated by NIP and NIL direct programming and RA patterning, we performed bulk mRNA sequencing at differentiation day 10. We sorted for VAChT-tdTomato negative and positive iCrMNs and iSpMNs and included iNeurog2/1 direct-programmed iNs to measure neuronal fate conversion and motor neuron differences in a principal component analysis (PCA) (**Figure 3a**)^31^. The PCA shows that neurons separate from iPSCs along PC1, accounting for 83% of the variance. PC1 also separates iNeurog2/1s from motor neurons. iCrMNs and iSpMNs then separate along PC2, accounting for 7% of the variance, demonstrating that they represent two similar yet distinct cell fates as was observed through immunostaining for specific fate markers (**Figure 2e-f**). The VAChT-tdT negative and positive cells for iCrMNs and iSpMNs cluster together in large part, aside from two replicates of iSpMNs in which the VAChT-tdT negative populations cluster slightly further apart. Taking the bulk RNAseq data together with comparisons between VAChT antibody stainings and VAChT-tdT marker expression, we concluded that the VAChT-tdT reporter likely underreports the number of motor neurons in the culture. Thus, the RNA-seq analysis supports the conclusion that direct programming by either NIP or NIL induction plus patterning with RA is sufficient to generate a nearly pure motor neuron culture after 10 days of differentiation.

**Figure 3.**
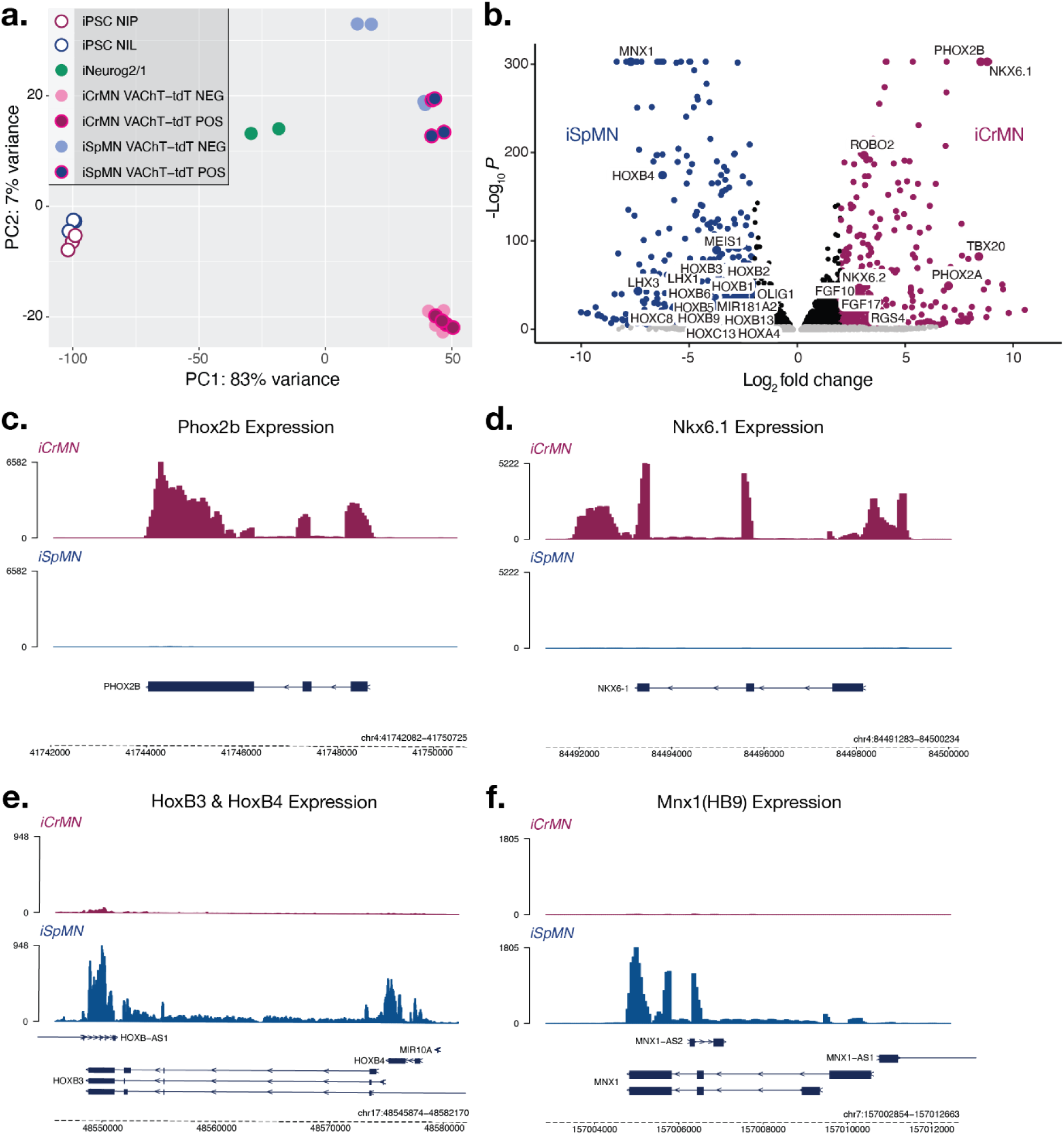
iCrMNs and iSpMNs differentially express genes associated with cranial and spinal motor neuron fate, respectively. (**a**) Principle Component Analysis (PCA) of iPSC NIP (white with red outline), iPSC NIL (white with blue outline), iNeurog2/1 (green), iCrMN VAChT- tdT NEG (light red), iCrMN VAChT-tdT POS (red with magenta outline), iSpMN VAChT- tdT NEG (light blue), and iSpMN VAChT-tdT POS (blue with magenta outline) normalized mRNA expression. (**b**) Volcano plot of genes differentially expressed in iSpMNs (blue, log_2_ fold change < -2, -log_10_P > 0.05) and iCrMNs (red, log_2_ fold change > 2, -log_10_P > 0.05). (**c**) BigWig normalized coverage reads for Phox2b in iCrMNs (red, above) and iSpMNs (blue, below). (**d**) BigWig normalized coverage reads for Nkx6.1 in iCrMNs (red, above) and iSpMNs (blue, below). (**e**) BigWig normalized coverage reads for HoxB3 & HoxB4 in iCrMNs (red, above) and iSpMNs (blue, below). (**f**) BigWig normalized coverage reads for Mnx1 (HB9) in iCrMNs (red, above) and iSpMNs (blue, below).

DESEQ2 differential expression analysis identified 7028 differentially expressed genes (**Figure 3b**)^34^. Among these, iCrMNs differentially express key cranial motor neuron fate markers such as *PHOX2B*, *NKX6.1*, *TBX20*, *FGF10*, *FGF17*, *RGS4*, *ROBO2*, and *NKX6.2*, markers which have been previously shown by *in-situ* hybridization and PAGODA analysis to have restricted expression in ALS-resistant cranial nerves^6,35–43^. iSpMNs differentially express key spinal motor neuron fate markers such as *MNX1* (*HB9*), *OLIG1*, *MEIS1*, and a range of Hox genes with *HOXB3* and *HOXB4* among the most differentially expressed. Of note, in previous studies in mouse embryonic stem cells, NIL expression alone was insufficient to turn on Hox gene expression^28^. Combining NIL induction with RA signaling generated mouse MNs with high Hox gene expression, a phenomenon we have now recapitulated in human iPSC-derived iSpMNs. Moreover, iSpMNs have significantly higher expression compared to iCrMNs of *MIR-181*, a microRNA recently found to be a prognostic biomarker of ALS^44^. We compared the gene expression coverage levels between iCrMN and iSpMN for a handful of select genes (**Figure 3c-f**). Expression of the cranial motor neuron markers *PHOX2B* and *NKX6.1* is exclusive to iCrMNs while expression of the spinal motor neuron markers *HOXB3*, *HOXB4*, and *MNX1* (HB9) is exclusive to iSpMNs. Note that although that HB9 immunocytochemistry revealed some signal in iCrMNs, RNA-seq did not capture any significant *MNX1* transcription in these neurons (**Figure 2f**, **Figure 3f**). Together, these findings demonstrate that direct programming by iNIP and iNIL plus differential patterning with RA is sufficient to rapidly generate human iPSC-derived cranial motor neuron and spinal motor fate with high efficiency.

### NIP and NIL direct programming generate ALS disease relevant CRISPR engineered mutant iCrMNs and iSpMNs

With the knowledge that NIP and NIL direct programming and RA patterning efficiently generate cranial and spinal motor neuron fate, respectively, we sought to utilize this technology to generate ALS disease relevant mutant lines. We used an available CRISPR engineered homozygous TDP43^M337V^ mutant iPSC line with a single copy doxycycline-inducible Neurog2. To this line, we performed the same transfection, selection, and clone induction and differentiation efficiency testing as we had for the VAChT-tdT lines (**Figure 1a**). Day 10 TDP43^M337V^ mutant iCrMNs and iSpMNs express the pan-neuronal markers β3-tubulin and MAP2 (**Figure 4a**). As was the case for the VAChT-tdT lines, TDP43^M337V^ iCrMNs express Nkx6.1 and Phox2b while TDP43^M337V^ iSpMNs express Hb9.

**Figure 4.**
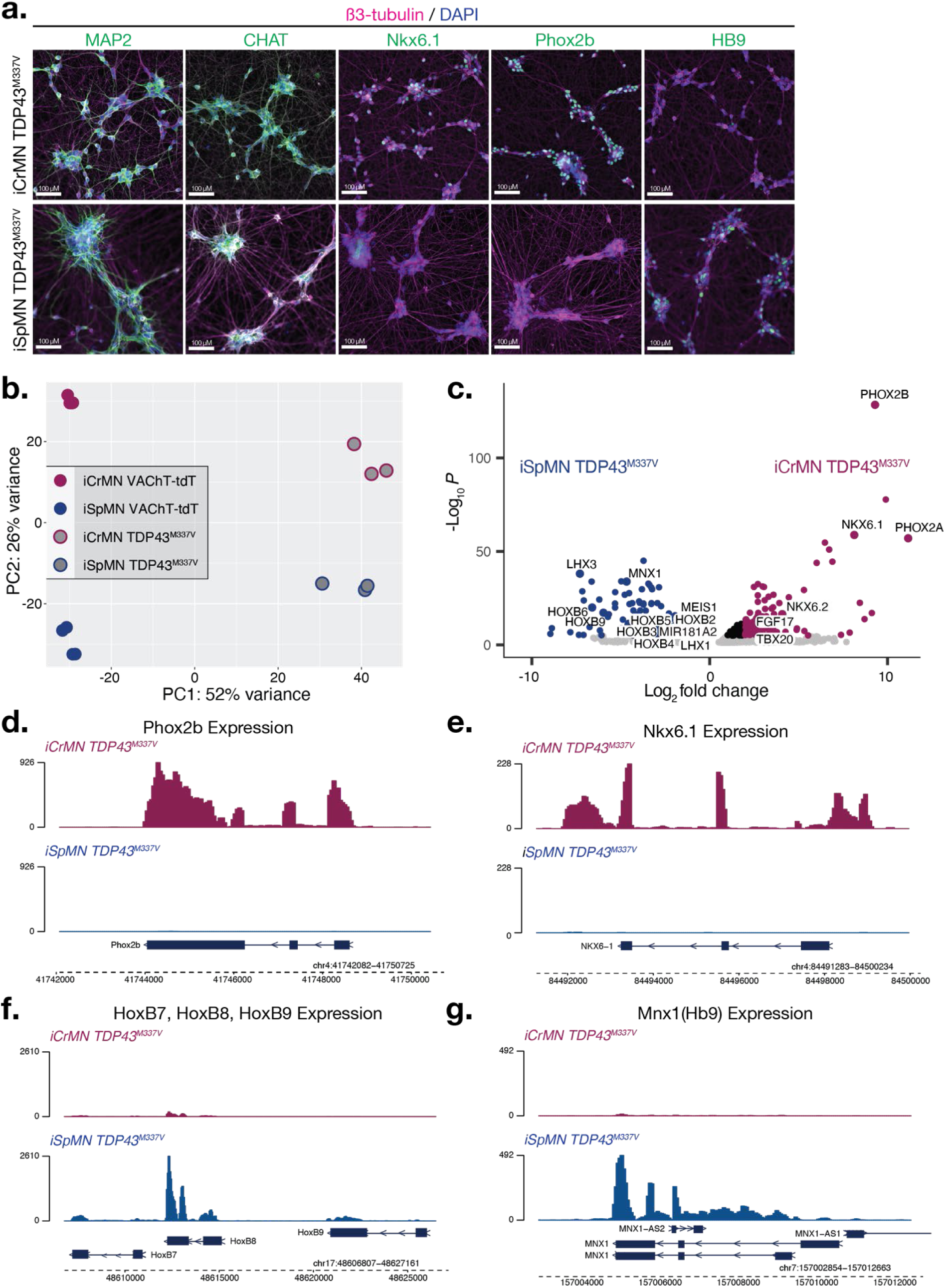
NIP and NIL direct programming and RA patterning differentiate CrMN and SpMN fate in an independent TDP43 mutant background. (**a**) Representative images for antibody staining of neuronal (MAP2, green, first from left), motor neuron (CHAT, green, second from left), cranial motor neuron (Nkx6.1, green, center; Phox2b, green, second from right), and spinal motor neuron (HB9, green, first from right) fate markers. Merged with DAPI (blue) and VAChT-tdT (magenta). Scale bars = 100 µm. (**b**) PCA of iCrMN VAChT-tdT (red), iSpMN VAChT-tdT (blue), iCrMN TDP43^M337V^ (grey with red outline), and iSpMN TDP43^M337V^ (grey with blue outline) normalized mRNA expression. (**c**) Volcano plot of genes differentially expressed in TDP43^M337V^ iSpMNs (blue, log_2_ fold change < -2, -log_10_P > 0.05) and iCrMNs (red, log_2_ fold change > 2, -log_10_P > 0.05). (**d**) BigWig normalized coverage reads for Phox2b in TDP43^M337V^ iCrMNs (red, above) and iSpMNs (blue, below). (**e**) BigWig normalized coverage reads for Nkx6.1 in TDP43^M337V^ iCrMNs (red, above) and iSpMNs (blue, below). (**f**) BigWig normalized coverage reads for HoxB7, HoxB8, & HoxB9 in TDP43^M337V^ iCrMNs (red, above) and iSpMNs (blue, below). (**g**) BigWig normalized coverage reads for Mnx1 (HB9) in TDP43^M337V^ iCrMNs (red, above) and iSpMNs (blue, below).

To compare the motor neuron fates generated in the TDP43 mutant background to that from VAChT-tdT lines, we used PCA analysis of day 10 iCrMNs and iSpMNs from both backgrounds (**Figure 4b**). PC1, explaining 52% of the variance among samples, defined the difference in cell line background, with TDP43 mutant lines separating away from the VAChT-tdT lines. PC2, explaining 26% of the variance, then defined motor neuron fate differences, with iCrMNs separating from iSpMNs. We performed differential expression analysis using DESEQ2 to identify differentially expressed genes between TDP43 mutant iCrMNs and iSpMNs (**Figure 4c**). Like the VAChT-tdT lines, iCrMNs differentially express key markers of cranial motor neuron fate such as *PHOX2B*, *NKX6.1*, *TBX20*, *FGF17*, and *NKX6.2*. iSpMNs differentially express key spinal motor neuron fate markers such as *MNX1* (*HB9*), *MEIS1*, and a range of Hox genes with *HOXB6* and *HOXB9* among the most significantly differentially expressed suggesting a more posterior spinal motor neuron fate than the VAChT-tdT iSpMNs. TDP43 mutant iSpMNs again have upregulated expression of the ALS prognostic biomarker *MIR-181*^44^. The cranial motor neuron markers Phox2b and Nkx6.1 are exclusive to TDP43 mutant iCrMNs, while expression of the spinal motor neuron markers *HOXB7*, *HOXB8*, *HOXB9*, and *MNX1* (*HB9*) is exclusive to TDP43 mutant iSpMNs (**Figure 4d-g**). Together, these findings demonstrate that in an entirely independent iPSC line carrying an ALS relevant mutation, NIP and NIL programming and RA patterning can efficiently generate cranial and spinal motor neuron fate, respectively.

### Human iCrMNs are more resistant than iSpMNs to chemically induced proteotoxic stress

With the ability to robustly and reproducibly generate human cranial and spinal motor neuron fates *in vitro*, we next sought to determine if NIP and NIL inducible motor neurons can recapitulate the differential sensitivity to ALS-related stress observed in patients^2–4^. ALS-sensitive spinal motor neurons are under proteostasis stress during ALS progression^11,45^. Following observations in mouse motor neurons, we hypothesized that under generalized proteotoxic stress, the superior ability of iCrMNs to maintain proteostasis would result in increased survival when compared to iSpMNs^7^. Thus, we compared the survival of iCrMNs and iSpMNs when treated with two well-described proteostatic stressors, cyclopiazonic acid (CPA) and tunicamycin (TM)^46,47^. CPA and Tunicamycin induce misfolded proteins by different mechanisms. CPA blocks endoplasmic reticulum Ca^2+^-ATPase, the Ca^2+^-pump in endoplasmic reticulum (ER) membrane that helps to maintain Ca^2+^ balance and ER homeostasis, and thus has a broad effect. Tunicamycin inhibits N-linked glycosylation and thus affects trans-membrane and secreted proteins. Notably, CPA was reported to accelerate the degeneration of SpMNs expressing hSOD1 G93A mutation, suggesting that CPA treatment may reveal ALS-relevant differential vulnerability between iCrMNs and iSpMNs^48^.

We treated iCrMNs and iSpMNs with CPA or TM on differentiation day 10 when terminal cell fates are established in culture. We then measured cell survival 18 days after treatment (differentiation day 28) with the colorimetric MTT assay (**Figure 5a**)^49^. iCrMNs were more resistant to treatment with CPA than iSpMNs across all tested concentrations (**Figure 5b**). To ensure that MTT measurement is a reliable proxy for survival, we utilized the VAChT-tdT reporter to image survival after CPA treatment and measured the binary area ratio over DAPI staining of nuclei after thresholding fluorescent intensity (**Supplemental Figure 3**). Although intrinsically less precise, measuring survival by imaging showed similar differential resistance of iCrMNs to CPA treatment compared to iSpMNs, demonstrating that MTT is a useful and high-throughput proxy method for measuring motor neuron survival. Treatment with TM revealed a similar phenotype, where iCrMNs were more resistant than iSpMNs to treatment with the three highest concentrations of TM tested (**Figure 5c**).

**Figure 5.**
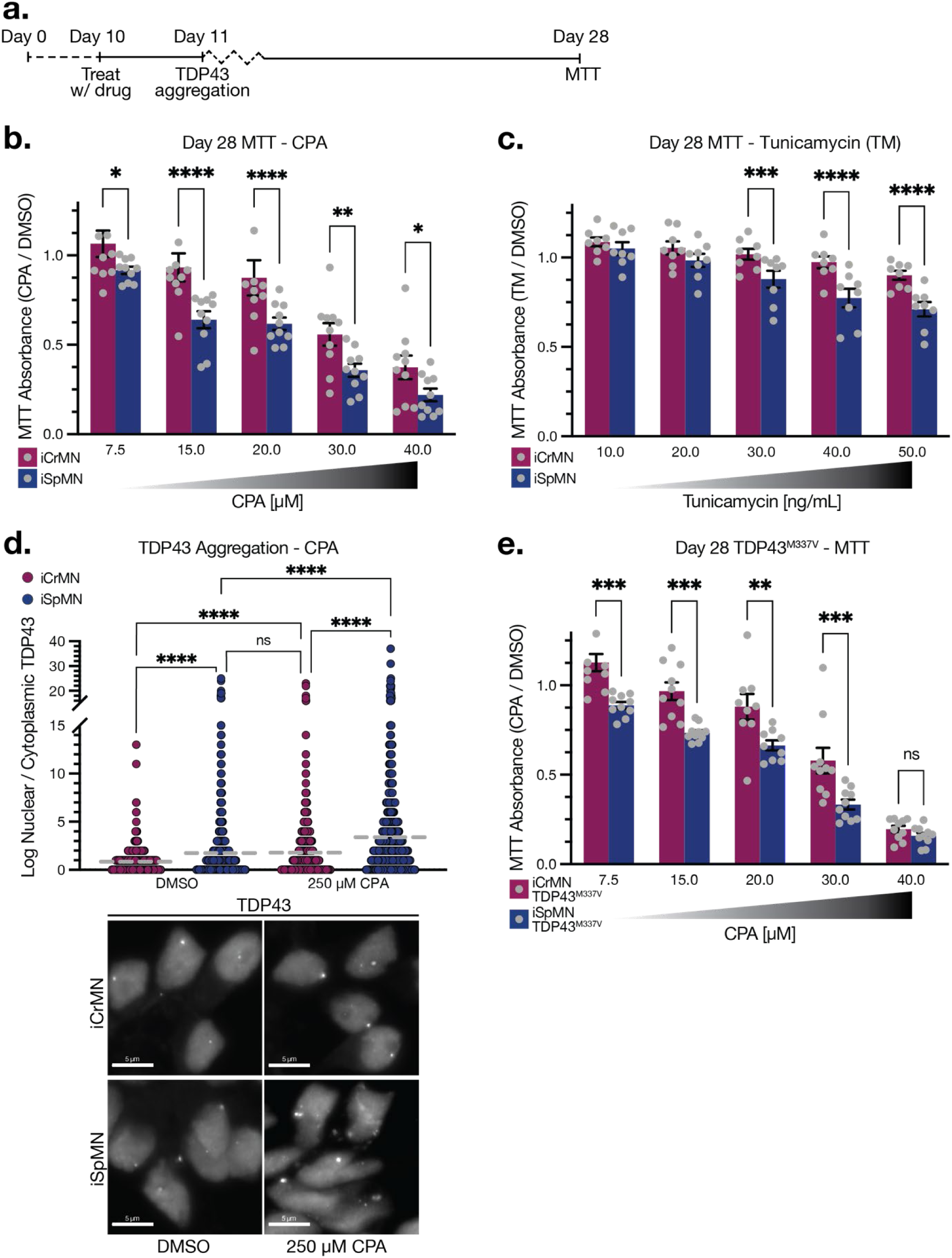
iCrMNs are more resistant than iSpMNs to proteotoxic stress. (**a**) Experimental outline: iCrMNs and iSpMNs are differentiated from day 0 to day 10, then treated with drug (CPA or Tunicamycin (TM)) on day 10. TDP43 aggregation is measured 24 hours after treatment. MTT cell viability is measured 18 days after treatment. (**b**) MTT measurement 18 days post treatment with CPA. iCrMNs are more resistant than iSpMNs to CPA treatment (n = 8-10, mean ± SEM, two-way ANOVA followed by Šídák’s multiple comparisons test). (**c**) MTT measurement 18 days post treatment with TM. iCrMNs are more resistant than iSpMNs to TM treatment (n = 8, mean ± SEM, two-way ANOVA followed by Šídák’s multiple comparisons test). (**d**) TDP43 aggregates per cell measured 24 hours post treatment with 250 µM CPA. iSpMNs accumulate more aggregates per cell than iCrMNs after proteotoxic stress (n = 3 biological replicates, >400 neurons per replicate, mean, Kruskal-Wallis Dunn’s test of multiple comparisons). Representative images below for TDP43 aggregation in DMSO (left) and 250 µM CPA (right) treated iCrMNs (top) and iSpMNs (bottom). Scale bar = 5 µm. (**e**) MTT measurement 18 days post treatment with TM. TDP43^M337V^ iCrMNs are more resistant than TDP43^M337V^ iSpMNs to CPA treatment (n = 8-10, mean ± SEM, two-way ANOVA followed by Šídák’s multiple comparisons test). ns = not significant, *p<0.0332, **p<0.0021, ***p<0.0002, ****p<0.0001.

We next wanted to determine if proteotoxic stress leads to ALS-related cellular phenotypes. TDP43 aggregate accumulation is a key hallmark of ALS progression across both familial and sporadic cases^13^. We treated iCrMNs and iSpMNs with a high CPA concentration (250 µM) for 24 hours and quantified TDP43 aggregate foci per cell (**Figure 5d**). iSpMNs contain more TDP43 aggregates per cell compared to iCrMNs (iCrMN DMSO 0.85 ± 1.08, iSpMN DMSO 1.75 ± 2.87, p < 0.0001). Treatment with CPA increases the number of TDP43 aggregates per cell in both iCrMNs and iSpMNs (iCrMN CPA 1.78 ± 2.62, iSpMN CPA 3.39 ± 4.14, p < 0.0001). Thus, the basal culture stress and CPA-induced stress led to more TDP43 aggregation in ALS-sensitive iSpMNs than iCrMNs.

The TDP43^M337V^ mutant iCrMNs and iSpMNs enabled us to ask whether the mutation accelerates cell death, thus eliminating all stress-sensitive motor neurons, or whether there are still neurons that would progressively die with additional stress. We treated TDP43^M337V^ mutant iCrMNs and iSpMNs with increasing concentrations of CPA on day 10 and measured cytotoxicity by MTT assay 18 days later on day 28 (**Figure 5e**). iCrMNs were again consistently significantly more resistant than iSpMNs across all tested concentrations of CPA. The similar response of iCrMNs and iSpMNs to CPA treatment between the VAChT-tdT and TDP43^M337V^ lines indicated that the TDP43 mutant background allows for motor neuron fate differentiation which maintains differentially susceptibility to stress within the timescale of the experiment. Similar to *in vivo* conditions or *in vitro* directed differentiation, direct programming of motor neurons carrying ALS-inducing mutations initially generates a stress-sensitive population^7^. Together, these results reflect the ability of directly programmed and patterned iCrMNs and iSpMNs to model *in vitro* the differential sensitivity to ALS observed in *in vivo* models and patients.

### Human iCrMNs rely on proteasome function to maintain proteostasis

The ubiquitin-proteasome system is the predominant pathway cells use to degrade proteins and maintain proteostasis via degradation^50^. Our previous study showed that mouse ESC derived iCrMNs neurons rely on a higher proteasome activity than iSpMNs to maintain proteostasis under stress. However, whether this difference extends to human CrMN biology is unknown. We hypothesized that human iCrMNs have the same reliance on proteasome function and thus would be more sensitive than iSpMNs to proteasome inhibition.

Cytoplasmic TDP43 mislocalization was previously shown to occur in iPSC derived spinal motor neurons after proteasome inhibition by MG132^20^. To determine if TDP43 localization is differentially affected in iCrMNs and iSpMNs after proteasome inhibition, we treated both cell types with high concentrations of MG132 or Epoxomicin (Epo), a different proteasome inhibitor, and measured TDP43 nuclear vs. cytoplasmic localization (**Figure 6a**). To measure nuclear versus cytoplasmic localization, we used DAPI to segment the nuclear TDP43 signal and measured the ratio of nuclear over cytoplasmic TDP43 signal per cell using CellProfiler (**Supplemental Figure 4**)^51^. iCrMNs retain nuclear TDP43 localization at higher rates compared to iSpMNs in control conditions (MG132: iCrMN DMSO 3.25 ± 1.99, iSpMN DMSO 3.05 ± 1.93, non-log transformed, p = 0.0001; Epo: iCrMN DMSO 5.66 ± 1.88, iSpMN DMSO 4.29 ± 2.00, non-log transformed, p = 0.0001) (**Figure 6b-c**). Treatment with 1 µM MG132 decreased the nuclear localization in both cell types, with iCrMNs mislocalizing TDP43 to the cytoplasm more than iSpMNs (iCrMN MG132 1.0 µM 1.57 ± 1.45, iSpMN MG132 1.0 µM 2.10 ± 1.54) (**Figure 6b**). The effect was exacerbated by 1.5 µM MG132 treatment (iCrMN MG132 1.5 µM 1.02 ± 1.50, iSpMN MG132 1.5 µM 1.54 ± 1.63, non-log transformed) (**Figure 6b**). Treatment with Epo decreased nuclear TDP43 localization only in iCrMNs (iCrMN Epo. 400 nM 3.38 ± 1.81, iSpMN Epo. 400 nM 3.95 ± 1.97, iCrMN Epo. 600 nM 2.85 ± 1.88, iSpMN Epo. 600 nM 4.76 ± 1.80, non-log transformed) (**Figure 6c**).

**Figure 6.**
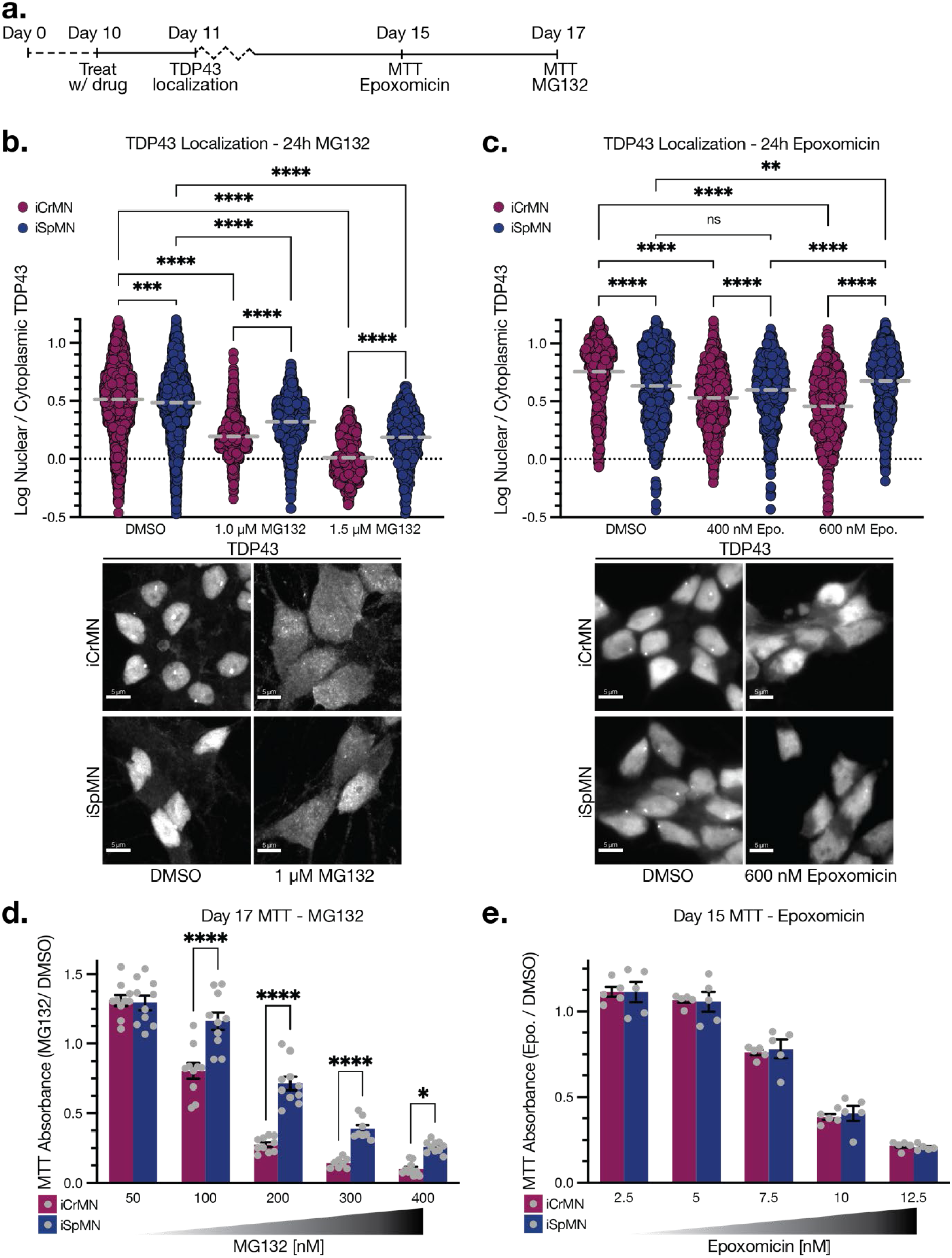
iCrMNs rely on the proteasome to maintain proteostasis. (**a**) Experimental outline: iCrMNs and iSpMNs are differentiated from day 0 to day 10, then treated with drug (MG132 or Epoxomicin) on day 10. TDP43 nuclear versus cytoplasmic localization is measured 24 hours after treatment. MTT cell viability is measured 5 or 7 days after treatment for Epoxomicin or MG132, respectively. (**b**) iCrMNs rely more on the proteasome to prevent TDP43 nuclear depletion by MG132 (n = 3 biological replicates, >400 neurons per replicate, mean, one-way ANOVA followed by Tukey’s multiple comparisons test). Representative images below for TDP43 localization in DMSO (left) and 1 µM MG132 (right) treated iCrMNs (top) and iSpMNs (bottom). (**c**) iCrMNs rely more on the proteasome to prevent TDP43 nuclear depletion by Epoxomicin (n = 3 biological replicates, >400 neurons per replicate, mean, one-way ANOVA followed by Tukey’s multiple comparisons test). Representative images below for TDP43 localization in DMSO (left) and 600 nM Epoxomicin (right) treated iCrMNs (top) and iSpMNs (bottom). (**d**) MTT measurement 7 days post treatment with MG132. iCrMNs are less resistant than iSpMNs to MG132 treatment (n = 10, mean ± SEM, two-way ANOVA followed by Šídák’s multiple comparisons test). (**e**) MTT measurement 5 days post treatment with Epoxomicin. iCrMNs and iSpMNs are equally sensitive to Epoxomicin treatment (n = 5, mean ± SEM, two-way ANOVA followed by Šídák’s multiple comparisons test). Scale bars = 5 µm. ns = not significant, *p<0.0332, **p<0.0021, ***p<0.0002, ****p<0.0001.

To test if proteasome inhibition would affect long-term motor neuron survival, we treated iCrMNs and iSpMNs with MG132 or Epo on day 10 and measured MTT 7 or 5 days later, respectively (**Figure 6a**). In response to MG132, iCrMNs were more sensitive than iSpMNs at the four highest concentrations tested (**Figure 6d**). Epo treatment was equally toxic for both iCrMNs and iSpMNs and MTT had to be measured two days earlier due to the increased toxicity for both cell types (**Figure 6e**). The long-term toxicity differences might be due to their different molecular action. MG132 is a synthetic peptide aldehyde that reversibly binds to the β5 subunit of the proteasome. Epoxomicin, on the other hand, covalently binds β7 and β10 proteasome subunits^52–55^. Together, these results suggest that mouse and human iCrMNs rely on proteasome function to prevent cellular vulnerability to ALS progression and maintain proteostasis.

### Cell type-specific splicing alterations in long-term TDP43 mutant cultures

In most ALS cases, TDP43 is depleted from the nucleus and accumulates in the cytoplasm of affected motor neurons^13,56^. To assess if homozygous TDP43^M337V^ mutation has an observable and differential effect on iCrMNs and iSpMNs, we stained for TDP43 to measure nuclear versus cytoplasmic localization at differentiation day 10 and day 30 in culture (**Figure 7a**). Day 10 TDP43^M337V^ iCrMNs retain higher rates of nuclear TDP43 localization compared to iSpMNs (iCrMN 4.92 ± 1.63, iSpMN 4.14 ± 1.61, non-log transformed, p < 0.0001) (**Figure 7b**). By day 30 there is no iCrMN reduction in TDP43 while iSpMNs further mislocalized TDP43 from the nucleus (iCrMN 5.07 ± 1.63, iSpMN 3.07 ± 1.89, non-log transformed, p < 0.0001). Thus, NIP and NIL induced motor neurons model a key feature of ALS disease progression where affected neurons mislocalize TDP43 in the cytoplasm with time.

**Figure 7.**
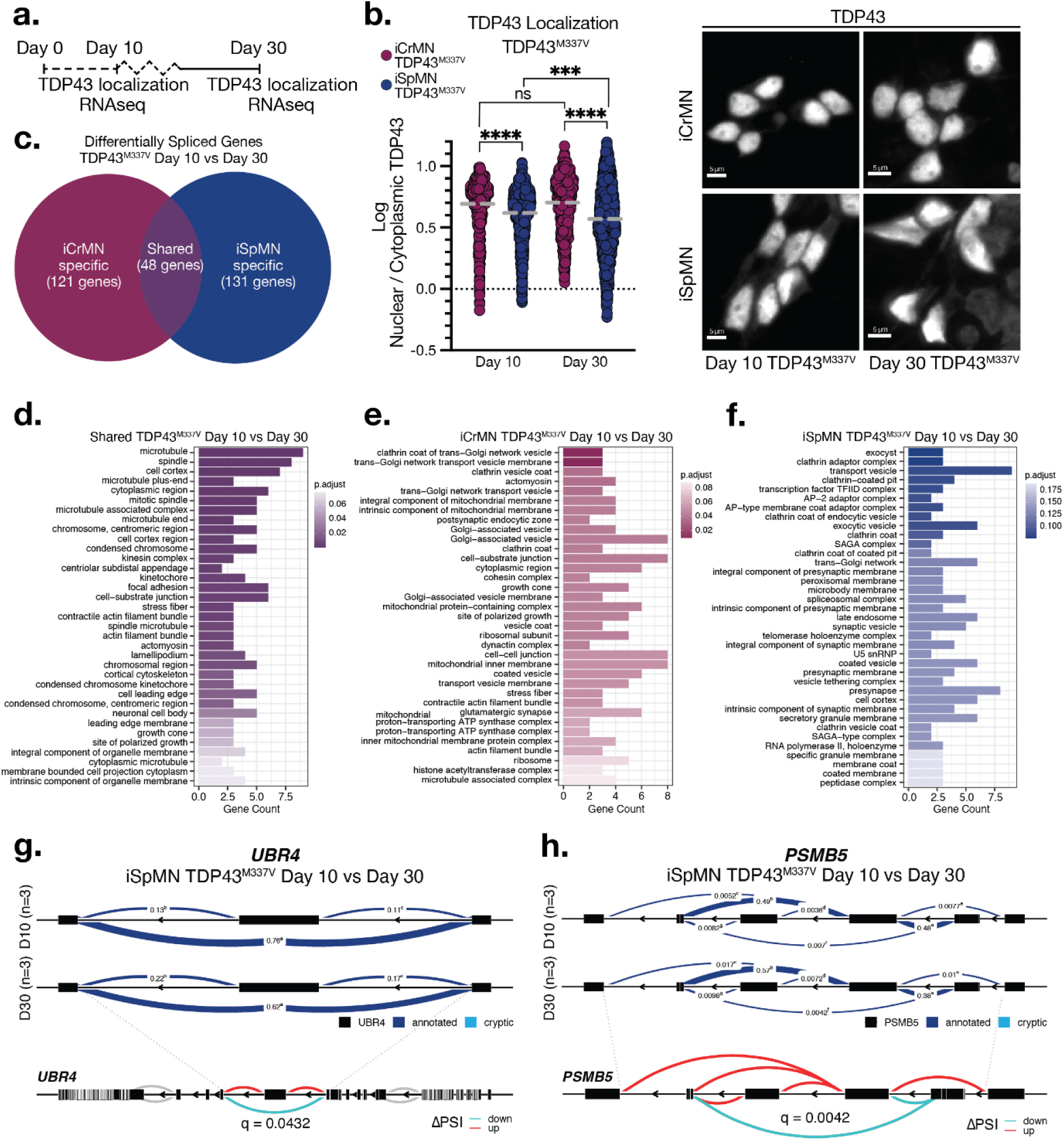
Cell type specific TDP43 localization and splicing changes over time in TDP43^M337V^ iCrMNs and iSpMNs. (**a**) Experimental outline: TDP43^M337V^ iCrMNs and iSpMNs are differentiated from day 0 to day 10 and fixed for TDP43 antibody staining and RNAseq on day 10 and day 30. (**b**) TDP43^M337V^ iCrMNs resist nuclear depletion of TDP43 compared to TDP43^M337V^ iSpMNs over time (n = 3 biological replicates, >400 neurons per replicate, mean, one-way ANOVA followed by Tukey’s multiple comparisons test). Representative images right for TDP43 localization in day 10 (left) and day 30 (right) treated iCrMNs (top) and iSpMN (bottom). Scale bar = 5 µm. (**c**) Day 10 versus day 30 differentially spliced genes (DSG) specific to TDP43^M337V^ iCrMNs (121 genes), iSpMNs (131 genes), and shared (48 genes) after the removal of DSGs between day 10 iCrMNs and iSpMNs. (**d**, **e**, & **f**) Gene ontology (GO) enriched biological functions for TDP43^M337V^ shared (**d**) iCrMN-specific (**e**) and iSpMN-specific (**f**) day 10 vs day 30 DSGs. (**g** & **h**) LeafCutter analysis of differentially spliced intron clusters in the proteostasis genes UBR4 (**g**) and PSMB5 (**h**). Exons, black boxes. Blue band thickness and inserted values represent proportion of spliced exon-exon pairs. Change in percent spliced in (ΔPSI).

TDP43 depletion from the nucleus leads to differentially spliced genes (DSG) and retention of intronic sequences creating so-called cryptic exons^18,19^. These cryptic exons can result in premature polyadenylation, stop codon truncation, and degrade transcripts, having a broad impact on cellular behavior and viability. TDP43 mutation and nuclear depletion of TDP43 have recently been shown to induce cryptic exon splicing in two genes, *UNC13A* and *STMN2*, both involved in neuronal synaptic function^20–23^. *STMN2* cryptic exon splicing was found after siRNA knockdown of TDP43 in iPSC-derived MNs and in neurons differentiated from ALS patient iPSC lines^20,21^. Cryptic exon splicing of *UNC13A* has also been observed in isolated neuronal nuclei from FTD-ALS patient frontal cortices^23^. However, it is unknown if TDP43 mutation differentially affects splicing in ALS sensitive SpMNs versus resistant CrMNs. To investigate the integrative, cumulative response of iCrMNs and iSpMNs to TDP43 localization over time, we used LeafCutter to analyze splicing alterations in TDP43^M337V^ MNs as the cells aged in culture^57^. We sequenced TDP43 mutant iCrMNs and iSpMNs at day 10 and at day 30 when we observed differential nuclear localization of TDP43 (**Figure 7a,b**). We performed pairwise comparisons of splicing differences in young day 10 versus aged day 30 TDP43^M337V^ iCrMNs and iSpMNs.

To focus specifically on the effect of TDP43 mutation induced splicing differences over time, we filtered out 11 DSGs between day 10 TDP34^M337V^ iCrMNs and iSpMNs. Although there may be a small fraction of differential splicing due to TDP43 localization differences already, day 10 differential splicing is mainly driven by cell fate differences and not the long-term effects of TDP43 mutation. After removing these DSGs, there were 169 DSGs in iCrMNs and 179 DSGs in iSpMNs after long term culture. Among these, 48 DSGs were shared, 121 DSGs were specific to iCrMNs, and 131 DSGs were specific to iSpMNs (**Figure 7c**). The relatively small number of shared DSGs indicates a cell-type specific response to TDP43 mutation over time in culture.

To investigate if these gene sets are associated with different biological functions, we compared the gene ontology (GO) enriched functional classes for DSGs specific to iCrMNs, iSpMNs, and shared in both cell fates (**Figure 7d-f**). Enriched pathways shared between iSpMN and iCrMN TDP43^M337V^ included microtubule, spindle, cell cortex, and microtubule, kinesin complex, stress fiber, actin filament bundle, and actomyosin, indicating splicing alterations in genes related to the cytoskeleton and active transport (all p < 0.05) (**Figure 7d**). GO analysis revealed iCrMN TDP43^M337V^ specific enrichment of pathways related to clathrin coat of trans-Golgi network vesicles, clathrin coated vesicles, actomyosin, mitochondrial membrane, and postsynaptic endocytic zone (all p < 0.05), suggesting altered splicing of genes involved in vesicle formation, mitochondrial function, and synaptic function (**Figure 7e**). Enriched pathways specific to iSpMN TDP43^M337V^ included exocyst, clathrin adaptor complex, and transport vesicle (all p < 0.1), suggesting altered splicing in genes related to vesicle exocytosis (**Figure 7f**). Although cryptic exon splicing of STMN2 has been observed in TDP43 mutant neurons, we did not observe any evidence of alternative splicing of *STMN2* in our TDP43^M337V^ iCrMNs or iSpMNs, nor of *UNC13A*.

Among the DSGs we found two genes spliced in iSpMNs but not in iCrMNs involved in the ubiquitin proteasome system: *UBR4* and *PSMB5*. Splicing of Ubiquitin Protein Ligase E3 Component N-Recognin 4 (*UBR4*), an E3-Ubiquitin ligase, is altered in day 30 versus day 10 TDP43^M337V^ iSpMNs, but not iCrMNs (q = 0.0432, ΔPSI = -0.137) (**Figure 7g, Supplemental Figure 5a**). TDP43 mutation over time leads to increased retention of an exon between exons 49 and 50 of the canonical transcript corresponding to exon 23 of the putative alternatively spliced isoform *UBR4-205* (Uniprot ID: A0A0A0MSW0). Of note, mutations to UBR4 are associated with episodic ataxia in patients^58,59^. *PSMB5* splicing is also altered in day 30 versus day 10 TDP43^M337V^ iSpMNs, but not iCrMNs (q = 0.0042, ΔPSI = -0.103) (**Figure 7h, Supplemental Figure 5b**). TDP43 mutation over time leads to increased retention of exons 1, 4, and 6 of *PSMB5*. Retention of these exons would likely lead to increased expression of alternatively spliced isoforms of *PSMB5-203*, *-205*, and *-206* (Uniprot IDs: P28074-3, P28074-2, and H0YJM8) at the expense of the canonical isoform *PSMB5-202* (Uniprot ID: P28074-1). Mouse SpMNs contain lower core proteasome protein levels (including PSMB5) than CrMNs and mouse and human SpMNs are more vulnerable to proteostatic stress than CrMNs^7^. Although the functions of these non-canonical differentially spliced isoforms remain unknown, decreased expression of the canonical isoform PSMB5 may further sensitize SpMNs to proteostatic stress.

### TDP43 nuclear depletion by proteasome inhibition induces cell-type specific splicing defects

To our knowledge, splicing defects linked to nuclear TDP43 loss have never been directly compared between ALS sensitive SpMNs and resistant CrMNs. Therefore, splicing differences could result from differences in TDP43 mislocalization between motor neuron types as well as cell-type specific responses to TDP43 mislocalization. Cryptic exon splicing of *STMN2* was shown after MG132 induced nuclear loss of TDP43 expression, linking proteasome function to TDP43 localization and RNA splicing^20^. Both the proteasome inhibitors MG132 and carfilzomib induce splicing changes^60,61^. Intron retention in SpMNs in two genes, *UNC13A* and *STMN2*, has recently been shown to be induced by TDP43 depletion from the nucleus^20–23^. Thus, we treated iCrMNs and iSpMNs with MG132 and asked if forced nuclear depletion of TDP43 leads to a similar or distinct set of DSGs 24 hours later (**Figure 8a**). Because of iCrMNs’ reliance on proteasome function and higher sensitivity compared to iSpMNs to MG132 treatment, we used MG132 concentrations which gave roughly equivalent levels of TDP43 mislocalization: 1.0 µM for iCrMNs and 1.5 µM for iSpMNs (**Figure 6b**). Differential splicing analysis between MG132 treated iSpMNs and iCrMNs uncovered significant retention of an intron skipping exons 37 and 38 in *UNC13A* (q = 0.00748, change in percent spliced in (ΔPSI) = 0.786) in MG132 treated iSpMNs but not in treated iCrMNs (**Figure 8b, Supplemental Figure 7a**). This event is distinct from the cryptic exon splicing observed between exons 20 and 21 recently reported^22,23^. Analysis with LeafCutter did not reveal any splicing differences in *STMN2* after MG132 treatment in either iCrMNs or iSpMNs. However, we observed expression of the reported cryptic exon sequence between exons 1 and 2 of *STMN2* after MG132 treatment of both iCrMNs and iSpMNs, confirming that nuclear TDP43 loss in NIP and NIL direct programmed MNs results in previously observed splicing defects (**Supplemental Figure 7b**)^20,21^.

**Figure 8.**
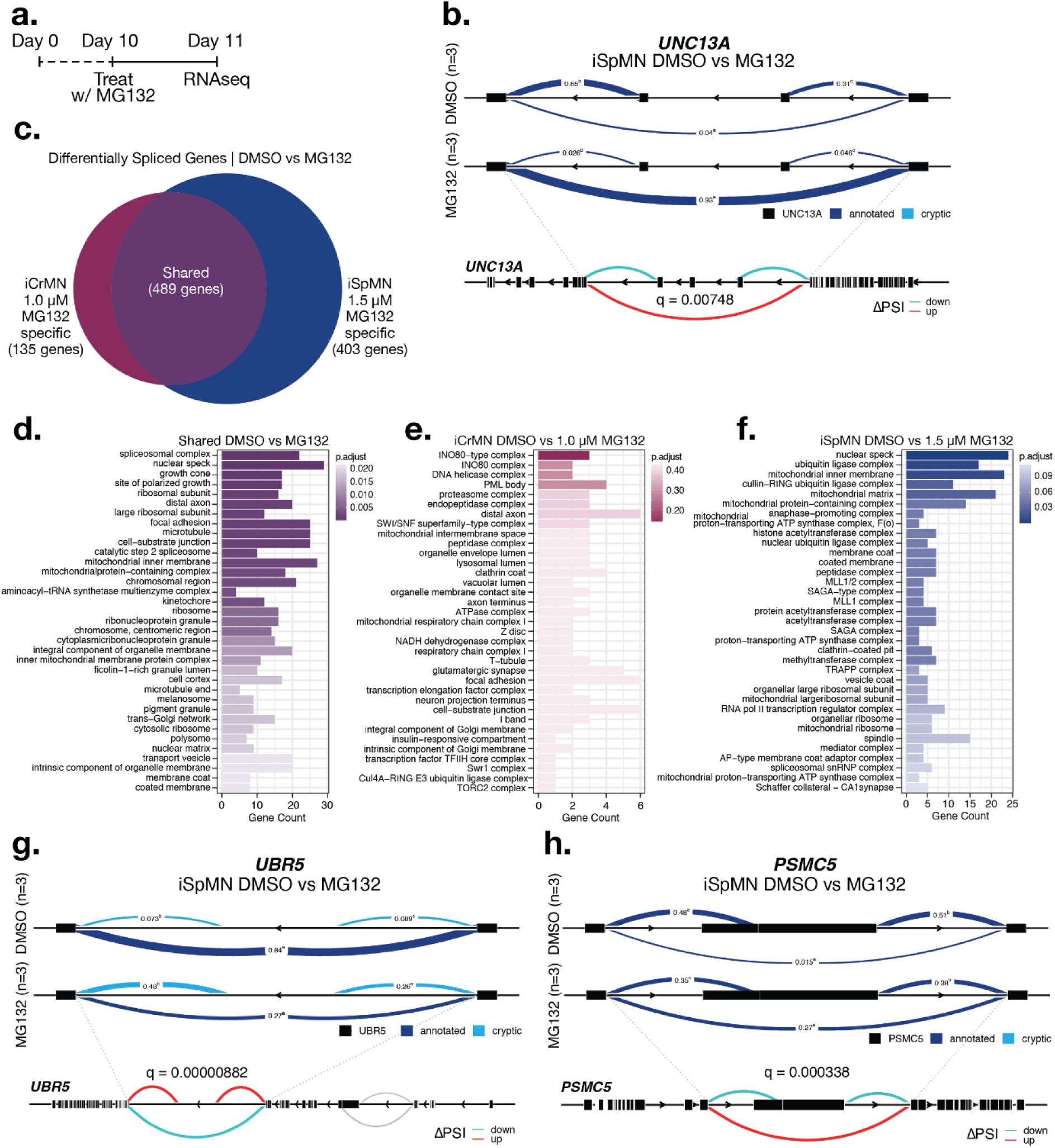
Cell type specific splicing changes after MG132 induced TDP43 nuclear depletion in iCrMNs and iSpMNs. (**a**) Experimental outline: iCrMNs and iSpMNs are differentiated from day 0 to day 10 and treated with 1.0 µM (iCrMN) or 1.5 µM (iSpMN) MG132 on day 10 and collected 24 hours late for RNAseq. (**b**) LeafCutter analysis of differentially spliced intron clusters in neuronal synaptic gene UNC13A. Exons, black boxes. Blue band thickness and inserted values represent proportion of spliced exon-exon pairs. Change in percent spliced in (ΔPSI). (**c**) DMSO versus MG132 DSGs specific to iCrMNs (135 genes), iSpMNs (403 genes), and shared (489 genes) after the removal of DSGs between DMSO treated iCrMNs and iSpMNs. (**d**, **e**, & **f**) Gene ontology (GO) enriched biological functions for shared (**d**) iCrMN-specific (**e**) and iSpMN-specific (**f**) DMSO vs MG132 DSGs. (**g** & **h**) LeafCutter analysis of differentially spliced intron clusters in the proteostasis genes UBR5 (**g**) and PSMC5 (**h**). Exons, black boxes. Blue band thickness and inserted values represent proportion of spliced exon-exon pairs. Change in percent spliced in (ΔPSI).

To understand how both neuron types respond to MG132-induced TDP43 nuclear depletion, we performed differential splicing analysis between DMSO and MG132 treatment in both cell types. In order to focus specifically on the effect of MG132, we filtered out 111 DSGs between DMSO treated iCrMNs and iSpMNs. After MG132 treatment 624 genes in iCrMNs and 892 genes in iSpMNs were differentially spliced. Of those, 489 were shared,135 were specific to iCrMNs, and 403 genes were specific to iSpMNs (**Figure 8c**). Since the levels of TDP43 nuclear depletion are roughly equivalent between the cell types at the chosen MG132 concentrations, the greater number of DSGs in iSpMNs suggests a much higher susceptibility to splicing defects compared to iCrMNs. iCrMNs, by comparison, seem to cope better in terms of TDP43 dependent splicing than iSpMNs to nuclear loss of TDP43.

We then asked if the two motor neuron types differentially spliced genes associated with different biological functions. We compared the GO enriched functional classes for DSGs specific to iCrMNs, iSpMNs, and shared in both cell fates (**Figure 8d-f**). Enriched pathways shared between iSpMN and iCrMN included spliceosomal complex, nuclear speck, growth cone, ribosomal subunit, distal axon, and focal adhesion, indicating splicing defects in a wide range of cellular processes including splicing itself (all p < 0.05) (**Figure 8d**). The small number of DSGs in MG132-treated iCrMNs meant that none of the GO terms were statistically significantly enriched. However, the analysis suggested DSGs were associated with the INO80 complex and DNA helicase complex (**Figure 8e**). Enriched pathways specific to iSpMN included nuclear speck, ubiquitin-ligase complex, mitochondrial inner membrane, and cullin-RING ubiquitin ligase complex (all p < 0.05), suggesting splicing defects in genes related to splicing itself as well as the ubiquitin-proteasome system and mitochondrial function (**Figure 8f**). To measure any effect on alternative splicing due to lower MG132 concentration used in iCrMNs, we also included data of iCrMNs treated with 1.5 µM MG132. 1.5 µM MG132 treatment compared to DMSO in iCrMNs led to 27 more DSGs compared to 1.0 µM treatment, with 20 additional genes specific to iCrMNs and 7 additional genes shared with iSpMNs (**Supplemental Figure 6a,b**). The greater number of DSGs after higher MG132 treatment correlates with increased nuclear depletion of TDP43 in iCrMNs at the same concentration (**Figure 6b**). GO enrichment was similar to treatment of iCrMNs with 1.0 µM MG132 (**Supplemental Figure 6c,d,e**).

### Altered splicing of proteostasis and spliceosomal complex related genes in iSpMNs

Among the DSGs in iSpMNs after MG132 treatment, we found genes related to the ubiquitin proteasome system (**Figure 9a**). Among these were *UBR5* and *PSMC5*. Ubiquitin Protein Ligase E3 Component N-Recognin 5 (*UBR5*) splicing is altered after MG132 treatment in iSpMNs but not iCrMNs (q = 8.82e-06, ΔPSI = -0.573) (**Figure 8g**, **Supplemental Figure 7c**). A cryptic exon inclusion between canonical exons 22 and 23 was observed only in MG132 treated iSpMNs. Sequence level inspection showed that a stop-codon truncation would occur across all three possible reading frames of the cryptic exon inclusion. UBR5 depletion in ALS and Huntington’s Disease patient iPSCs has been shown to induce aggregate formation, with UBR5 overexpression resulting in polyubiquitination and thus deceased levels and aggregation of mutant HTT^62^. Several subunits of the proteasome are also differentially spliced specifically in iSpMNs after MG132 treatment (**Figure 9a**). The 19S regulatory subunit *PSMC5* showed decreased inclusion of exon 5 in MG132 treated iSpMNs (q = 3.38e-04, ΔPSI = 0.258) (**Figure 8h, Supplemental Figure 7d**). Loss of exon 5 is found in the alternatively spliced isoform *PSMC5-213* (Uniprot ID: J3QQM). Separate intron retention events in *PSMC5* have been previously observed in a comparison between healthy and C9ORF72 mutant ALS patient samples, among splicing changes across multiple other proteasome subunits^63^. *UBR5* as well as subunits of the proteasome thus appear to be targets of disrupted splicing under ALS-related cellular stresses.

**Figure 9.**
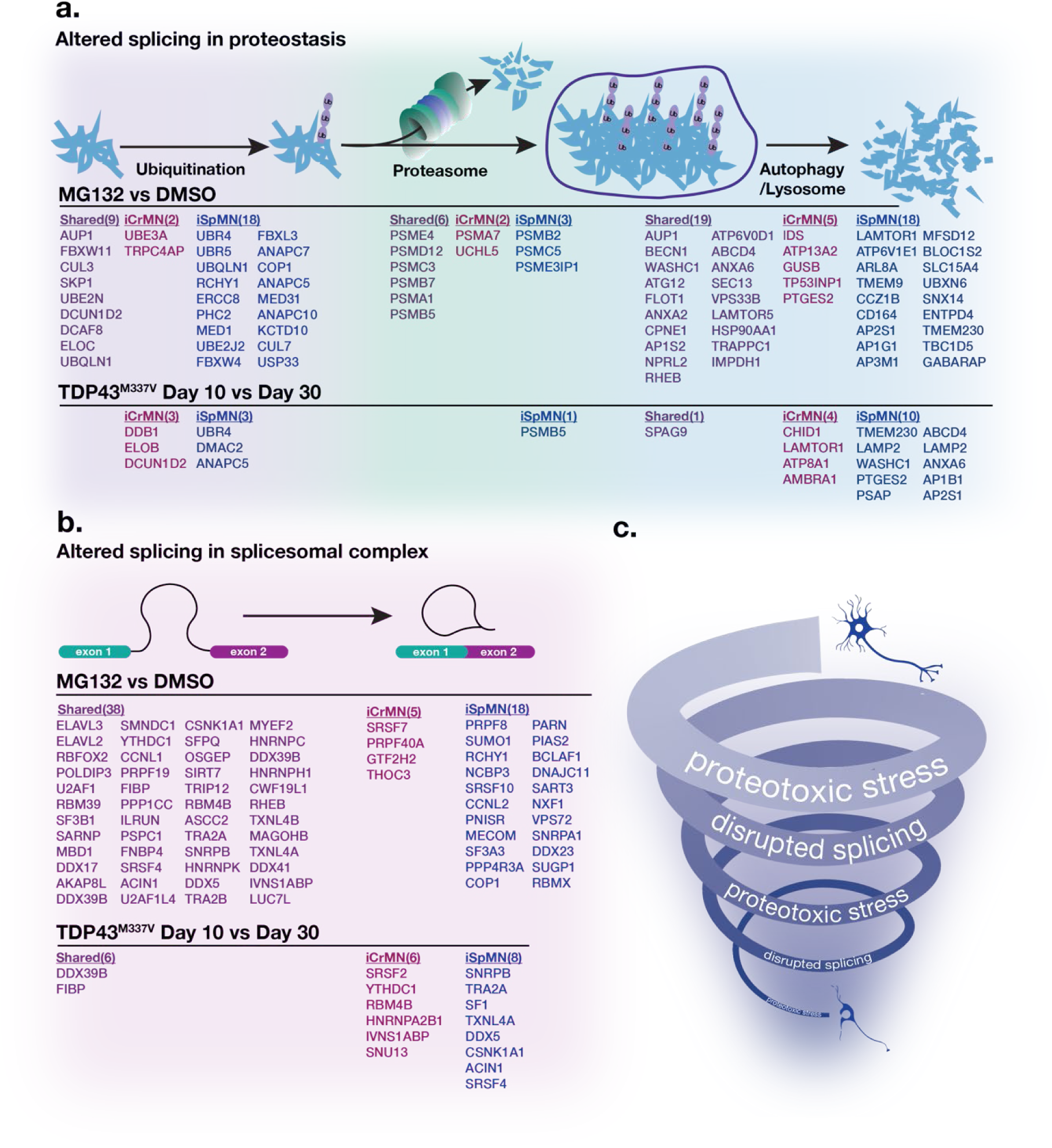
iSpMNs differentially splice more genes than iCrMNs related to proteostasis and splicing after MG132 treatment and overtime with TDP43 mutation. **(a)** Proteostasis pathway diagram with shared (violet), iCrMN-specific (red), and iSpMN-specific (blue) DSGs related to ubiquitination (left, pink background), proteasome (center, green background), and autophagy or lysome (right, blue background) after MG132 treatment (across middle) or over time with TDP43 mutation (across bottom). (**b**) RNA splicing diagram with shared (violet), iCrMN-specific (red), and iSpMN-specific (blue) DSGs related to splicesomal complex. (**c**) Sensitive SpMNs enter a downward spiral of proteotoxic stress and disrupted splicing leading to degeneration.

Strengthening the proteostatic stress and splicing connection, spliceosomal complex was the top GO enriched term in DSGs shared between iCrMNs and iSpMNs after MG132 treatment (**Figure 8d**). Among the shared DSGs involved in spliceosomal complex we found *ELAVL3*. TDP43 knockdown results in decreased expression and a cryptic exon event in *ELAVL3* in ALS patient derived MNs^20^. Additionally, ELAVL3 has recently been shown to be depleted from nuclei of sensitive neurons in ALS patient samples^64^. A cryptic exon inclusion between exons 3 and 4 of *ELAVL3* was observed in both iCrMNs and iSpMNs by LeafCutter analysis (iCrMN q = 1.43e-07, ΔPSI = -0.535; iSpMN q = 5.25e-07, ΔPSI = -0.529) (**Supplemental Figure 7e**). Read coverage plotting revealed clear expression of the cryptic exon in both cell types after MG132 treatment (**Supplemental Figure 7d**). Sequence level inspection showed that inclusion of this cryptic exon results in premature stop-codon truncation in all three reading frames, possibly explaining the decreased expression seen in MG132 versus DMSO treated iCrMNs and iSpMNs (**Supplemental Figure 7f**). Thus, TDP43 nuclear loss appears to lead to altered splicing of genes that are themselves involved in splicing, with the potential to disrupt splicing further.

We compared the number of DSGs after MG132 treatment exclusive to iCrMN, iSpMNs, or shared in both cell types belonging to GO terms related to ubiquitination, proteasome function, and autophagy or lysosomal function (**Figure 9a**) as well as spliceosomal complex (**Figure 9b**). There are 9 shared DSGs related to ubiquitination, with 2 specific to iCrMNs and 18 specific to iSpMNs. There are 6 shared differentially spliced components of the proteasome, with 2 specific to iCrMNs and 3 specific to iSpMNs. There are 19 shared DSGs related to autophagy or lysosomal function, with 5 specific to iCrMNs and 18 specific to iSpMNs. There are 38 shared DSGs related to spliceosomal complex, with 5 specific to iCrMNs and 18 specific to iSpMNs. We also looked at DSGs after prolonged TDP43 mutation and found a similar prevalence of iSpMN specific targets (Ubiqutination: 0 Shared, 3 iCrMN, 3 iSpMN; Proteasome: 0 Shared, 0 iCrMN, 1 iSpMN; Autophagy/Lysosome: 1 Shared, 4 iCrMN, 10 iSpMN; Spliceosomal complex: 2 Shared, 6 iCrMN, 8 iSpMN). Thus, although at lower numbers, the mild stress induced by the TDP43 mutation that causes degeneration over the course of years in patients induces differential splicing consequences within the span of days in these neurons.

Together, the comparison between iCrMNs and iSpMNS revealed a motor neuron type specific difference in their responses to ALS-related stresses. ALS-resistant motor neurons are more resistant in their ability to prevent mislocalization and precipitation of TDP43, SOD1, or p62, as we have shown before. Additionally, CrMNs also seem to be more resistant to the consequences of TDP43 mislocalization. Altered splicing of proteostasis-associated genes suggests that sensitive motor neurons engage in a downward spiral, wherein reduced capacity to maintain proteostasis leads to nuclear TDP43 reduction, resulting in a lower ability to prevent the accumulation of TDP43 and other misfolding proteins (**Figure 9c**).

## Discussion

Why some neurons die while others survive is a critical question in understanding the underlying pathology of various neurodegenerative diseases and uncovering effective therapies. In ALS, sensitive SpMNs progressively degenerate while a subset of resistant CrMNs survive. To investigate the nature of MN differential sensitivity, we developed an optimized rapid, efficient, and scalable protocol to directly program and positionally pattern iCrMNs and iSpMNs. However, a challenge in studying neurodegenerative diseases *in vitro* is modeling patient-relevant tissue complexity and long-term, late-stage degeneration in response to chronic low-level stress from single mutations. Complex, long-term tissue modeling is beginning to address these limitations through organoid, assembloid, and organ-on-chip models, which incorporate multiple cell types and even distinct tissues, as in a neuromuscular assembloid or cortical-blood-brain-barrier organ chip^65–68^. Because they introduce cell type interactions, these technologies hold significant therapeutic promise to model differential sensitivity to ALS and other neurodegenerative diseases. However, these technologies are usually labor-intensive and present significant complications to perform biochemical studies in individual cell types. On the other hand, the direct programming technology presented here enables the investigation of CrMN and SpMN intrinsic differences in response to ALS-related stress due to its high efficiency, allowing bulk biochemical and expression analyses without the need for cell sorting. Additionally, the rapid, monolayer differentiation protocol enables platform scalability conducive for high-throughput drug or gene-therapy screens.

Our previous work in *in vitro* mouse modeling of ALS revealed that CrMNs have a superior ability to maintain proteostasis due to a greater reliance on proteasome function than SpMNs. Mimicking the differential sensitivity observed in patients and *in vivo* and *in vitro* ALS models, programmed human iCrMNs are more resistant than iSpMNs to ALS-related chemically induced proteotoxic stress^2–6^. Human iCrMN resistance to ALS also appears to be due to their higher reliance on proteasome function to maintain proteostasis. Similar to mouse iCrMNs preventing SOD1 precipitation, human iCrMNs control TDP43 cytoplasmic accumulation. Disrupted proteostasis and ER stress play key roles in SpMN sensitivity to ALS^8–11,45,50,69–72^. Even in the relatively short time frame allowed by *in vitro* studies presented here, iSpMNs mislocalized mutant TDP43 over time, while iCrMNs retained nuclear TDP43. Thus, the ability to maintain a healthy proteome seems to be an intrinsic difference between ALS-resistant and sensitive MNs.

Despite the historical difficulties in accessing or differentiating subpopulations of ALS-resistant MNs, a limited number of studies have investigated differential sensitivity. Matrix metallopeptidase 9 (MMP-9) is differentially upregulated in ALS-vulnerable fast-fatigable α-SpMNs compared to resistant Onuf’s nucleus MNs^73^. In a hSOD1 G93A mouse model of ALS, MMP-9 reduction delayed muscle denervation of fast-fatigable α-SpMNs. Insulin-like growth factor 2 (IGF-2), on the other hand, is upregulated in resistant oculomotor neurons in mice, and muscular overexpression of IGF-2 extended the lifespan of hSOD1 G93A mice^74^. Targeted metabolomics across 17 ALS iPSC lines revealed increased arachidonic acid levels in differentiated SpMNs compared to ocular motor neurons, with pharmacological reduction reversing ALS-related phenotypes in SpMNs as well as *in vivo* fly and mouse ALS models^75^. To those processes potentially contributing to differential sensitivity, the results presented here add proteostasis and splicing. Thus, several different pathways and biological functions may underly differential sensitivity to ALS.

Many studies have investigated ALS-related disrupted splicing in sensitive SpMNs^18,20–22,56,63,76–81^. With forced nuclear depletion of TDP43 by MG132 treatment, the number of DSGs was nearly 3-fold higher in iSpMNs compared to iCrMNs, suggesting that SpMNs are differentially vulnerable to disrupted splicing. This is despite the fact that iSpMNs were treated with a higher concentration of MG132 than iCrMNs to achieve similar levels of TDP43 nuclear depletion. Thus, iCrMNs prevent TDP43 localization and resist splicing defects induced by its forced nuclear depletion. Among the biological functions of DSGs shared between iCrMNs and iSpMNs after TDP43 nuclear depletion, spliceosomal complex was the top category. Impaired splicing thus appears to lead to further, autoregulated impairment of splicing. Autoregulation of splicing has been shown wherein nuclear depletion of TDP43 leads to a cryptic exon event in the RBP HNRNPA1, which results in an aggregation prone variant thus further disrupting RNA splicing^82^. Additionally, ALS causing mutation to the RBP fused in sarcoma (FUS) leads to intron-retention events in other RBPs, including in FUS itself^79^. This splicing cascade is well exemplified by a cryptic exon inclusion in the RBP *ELAVL3* introducing a premature stop codon, which was also found previously^20^. Nuclear depletion of ELAVL3 has been found to precede and be more widespread than that of TDP43 in sensitive SpMNs from 40 different ALS patient samples^64^. While a large number are shared, the higher number of iSpMN-specific splicing related DSGs compared to iCrMN suggests that iSpMNs are particularly susceptible to autoregulation of splicing.

Additionally, DSGs with functions related to proteostasis maintenance were over-represented in iSpMNs compared to iCrMNs. More splicing differences were related to ubiquitination, proteasome function, and autophagy or lysosomal function in MG132 treated iSpMNs compared to iCrMNs. Cryptic exon splicing of the key autophagy gene *ATG4B* has been shown in ALS patient samples and after TDP43 knockdown^83^. Thus, an ALS-sensitive SpMN-specific downward spiral dynamic begins to emerge in which disrupted proteostasis, through proteasome inhibition, causes disrupted splicing and nuclear depletion of TDP43 and perhaps other RBPs. TDP43 nuclear depletion subsequently leads to autoregulated splicing defects in splicing-related genes and in genes related to proteostasis, which possibly further exacerbates splicing defects.

In summary, this work adds two possible mechanisms that allow some motor neurons to resist neurodegenerative stress. First, resistant neurons have a better ability to prevent protein accumulation, including aggregation of typical ALS proteins SOD1, p62, and TDP43. Second, resistant neurons have a greater capacity to resist splicing changes induced by TDP43 nuclear depletion. Thus, there is therapeutic potential in increasing the ability of ALS-sensitive to resist either proteotoxic stress or disrupted splicing, or both together. In the combined context comparing mouse and human motor neurons, proteasome function appears to play a potentially critical upstream role in differential sensitivity to ALS-related stress and resistance to disease progression. The fact that a large number of ALS causative familial mutations are in genes with functions related to proteostasis, RNA-binding, and splicing supports a mechanistic link between these processes^56^. There may be a critical threshold of resistance to the proteotoxic stress-disrupted splicing downward spiral which is higher in resistant CrMNs than sensitive SpMNs and that, once passed, leads to a rapid onset of degeneration late in life.

## Materials and Methods

### iPSC lines

The human iPSC VAChT-tdT line (from PMID: 32615233) was received from the Wichterle lab and the Stem Cell Core Facility at Columbia University Irving Medical Center. The human iPSC homozygous TDP43^M337V^ mutant line was generated by CRISPR-Cas9 engineering to introduce the M337V mutation to the *TARDBP* gene in a KOLF2.1J genetic background^84^. Integration of the NIP and NIL transcription factor cassettes was performed as previously described^30,85^. After Blasticidin selection, 32 clones per NIP and NIL were picked to wells of a Matrigel (Corning) coated 96 well plate and expanded. Clones were assessed for induction efficiency by Isl1 (DSHB 39.4D5) staining 2 days after doxycycline induction. Clones with high Isl1 induction efficiency were then differentiated for 10 days (protocol below) and stained for VAChT (Thermo Fisher Scientific, MA5-27662) or imaged for VAChT-tdT expression to assess MN differentiation efficiency. The most efficient clones were then expanded and maintained. NIP and NIL integration sites were mapped in selected clones by shotgun whole-genome sequencing using the MinION Nanopore platform in collaboration with FindGenomics. iPSCs were maintained in Essential 8 media (Gibco), supplemented with ROCK inhibitor (Enzo Life Sciences, Y-27632) for 24 hours after passaging or thawing, on Matrigel coated plates.

### iCrMN and iSpMN Differentiation

NIP and NIL iPSCs were split when cultures reached ∼70-80% confluence and plated at a density of 8000 cells/cm^2^ on Matrigel coated plates in Neurobasal-A (Gibco) media containing 1x Glutamine (Gibco), 1x N2 (Thermo Fisher Scientific), 1x B-27 (Thermo Fisher Scientific), 10 µM Y-27632, and 3 µM doxycycline. Media was changed every two days for 10 days. On day 2, media was changed to add 0.01 µM or 1.0 µM retinoic acid for iCrMN or iSpMN differentiation, respectively, and maintained throughout differentiation. On day 4, media was changed to remove Y-27632. On day 6, a half media change was performed to add anti-mitotic compounds, 4 µM 5-Fluoro-20-deoxyuridine (FDU) and 4 µM Uridine (Sigma). On day 8, a half media change was performed to add 200 ng/mL L-ascorbic acid (Sigma), 10 ng/mL BDNF, 10 ng/mL CNTF, 10 ng/mL GDNF (Peprotech). Day 10 was established as the end point for differentiation of terminal CrMN or SpMN fate.

### Immunofluorescence

Cells were fixed with 37°C 4% paraformaldehyde (Electron Microscopy Sciences) in 1X PBS for 15 minutes. Cells were incubated for 5 minutes in permeabilizing solution (0.2% Triton X (Sigma) in 1X PBS) then washed with 1X PBS. Cells were incubated for 1 hour in blocking solution (10% FBS (Sigma) in 1X PBS). Subsequently, cells were incubated with primary antibodies for 2 hours at room temperature or overnight at 4°C, washed in 1X PBS 3 times, and incubated with fluorescence-conjugated secondary antibodies for 1 hour at room temperature. The primary antibodies used were: Mouse anti-Tubb3, 1:200, BioLegend, MMS-435P; Chicken anti-MAP2, 1:5000, Abcam, ab5392; Mouse anti-VAChT, 1:500, Thermo Fisher Scientific, MA5-27662; Mouse anti-Phox2b, 1:100, Santa Cruz, sc-376997; Mouse anti-Nkx6.1, 1:100, DSHB, F55A10-c; Rabbit anti-HB9, 1:100, Millipore Sigma, ABN174; Rabbit anti-CHAT, 1:1000, Abcam, ab181023; Rabbit anti-TDP43, 1:250, ProteinTech, 10782-2-AP). The secondary antibodies used were: Alex Fluor Goat anti-(Mouse/Rabbit/Chicken) (488/568/647), 1:500 or 1:1000, Thermo Fisher Scientific. Cells were mounted in Fluoroshield with DAPI (Thermo Fisher Scientific). Tiled and representative images for quantification of cell fate markers were captured using a Nikon Perfect Focus Eclipse Ti live cell fluorescence microscope using the 10X and 20X objectives respectively. The Nikon Elements software was used to perform binary spot detection or thresholding for image quantification. Imaging for TDP43 aggregation and localization was performed using a Zeiss LSM 880 confocal microscope using the 63X oil immersion objective.

### RNA sequencing and analysis

RNA was extracted by using TRIzol LS Reagent (Life Technologies) and purified using the RNAeasy Mini Kit (Qiagen). Agilent High Sensitivity RNA Screentape (Agilent, 5067-5579) was used to check RNA integrity. A 500 ng quantity of RNA was used to prepare RNA-seq libraries and spiked-in with ERCC ExFold Spike-In mixes (Thermo Fisher Scientific, 4456739). RNA-seq libraries were prepared using a TruSeq Stranded mRNA Library Prep kit (Illumina, 20020594). Library size was verified using High Sensitivity DNA ScreenTape (Agilent, 5067-5584). The KAPA Library Amplification kit was used on a Roche LightCycler 480 for library quantification before pooling. The libraries were sequenced on an Illumina NovaSeq 6000 (SP, 100 cycles, single-end 100 bp) at the Genomics Core Facility at New York University. Fastq files obtained from RNA-seq were aligned to Ensembl and UCSC hg38 genomes using the splice-aware STAR (Spliced Transcripts Alignment to a Reference) aligner (v2.7.6a)^86^. Alignment files were indexed using samtools and converted to bigwig format using deeptools^87–89^. Mapped reads were assigned to gene annotated Ensembl and UCSC hg38 gene transfer files using the featureCounts function in subread (v2.0.1)^90,91^. Read counts were normalized using the ‘rlog’ or regularized log transformation in DESeq2 (v1.20.0)^34^. Transformed read counts were used as input features into the dimensionality reduction techniques PCA. Coverage plots for gene expression were generated using trackplot in bwtools^92^. Differential splicing and intron retention analysis was performed using LeafCutter^57^. GO enrichment analysis was performed using DOSE and EnrichPlot^93,94^.

### Cell survival and viability

Cell survival was measured by tile imaging direct fluorescence of the VAChT-tdTomato reporter in iCrMNs and iSpMNs differentiated in a 96 well plate and treated with cyclopiazonic acid (CPA, C1530-10MG, Sigma) on differentiation day 10. Imaging was performed using a 10X objective on a Nikon Perfect Focus Eclipse Ti live cell fluorescence microscope. Images were binary thresholded for fluorescent intensity and the binary area was measured per condition as a readout of cell survival. Cellular viability was measured in iCrMNs and iSpMNs differentiated in a 96 well plate and treated with CPA, tunicamycin (T7765-10MG, Sigma), or MG132 using a MTT 3-(4,5-dimethylthiazol-2-yl)-2,5-diphenyltetrazolium-based in vitro toxicology kit (TOX1-1KT, Sigma). 10 µl MTT solution was applied to cells at the end point of the experiment and incubated for 4 hours. 100 µl solubilization solution was added directly to wells and incubated overnight at 37°C. The MTT colorimetric assay was measured using a Tecan Spark Microplate Reader at an absorbance wavelength of 570 nm. Statistical analysis was performed using two-way ANOVA followed by Šídák’s multiple comparisons test.

### TDP43 Aggregation and Localization

Measurement of TDP43 aggregation was performed after 24-hour treatment of iCrMNs and iSpMNs with CPA (C1530-10MG, Sigma) followed by immunofluorescence and imaging using a Zeiss LSM 880 confocal microscope with a 63X oil immersion objective. 5 images were taken per condition per 3 biological replicates. Image analysis and quantification of TDP43 aggregates per cell was performed using CellProfiler^51^. Statistical analysis was performed using a Kruskal-Wallis test followed by a Dunn’s test of multiple comparisons. Measurement of TDP43 localization was performed on untreated TDP43^M337V^ iCrMNs and iSpMNs or VAChT-tdT iCrMNs and iSpMNs treated with MG132 (474790-1MG, Sigma) or Epoxomicin (324801-50UG, Sigma) followed by immunofluorescence and imaging using a Zeiss LSM 880 confocal microscope with a 63X oil immersion objective. 5 images were taken per condition per 3 biological replicates. To measure nuclear versus cytoplasmic localization, DAPI was used to segment the nuclear TDP43 signal and the ratio of nuclear over cytoplasmic TDP43 signal per cell was measured using CellProfiler^51^. Statistical analysis was performed using ANOVA followed by a Tukey’s multiple comparisons test.

---

**Supplemental Figure 1.**
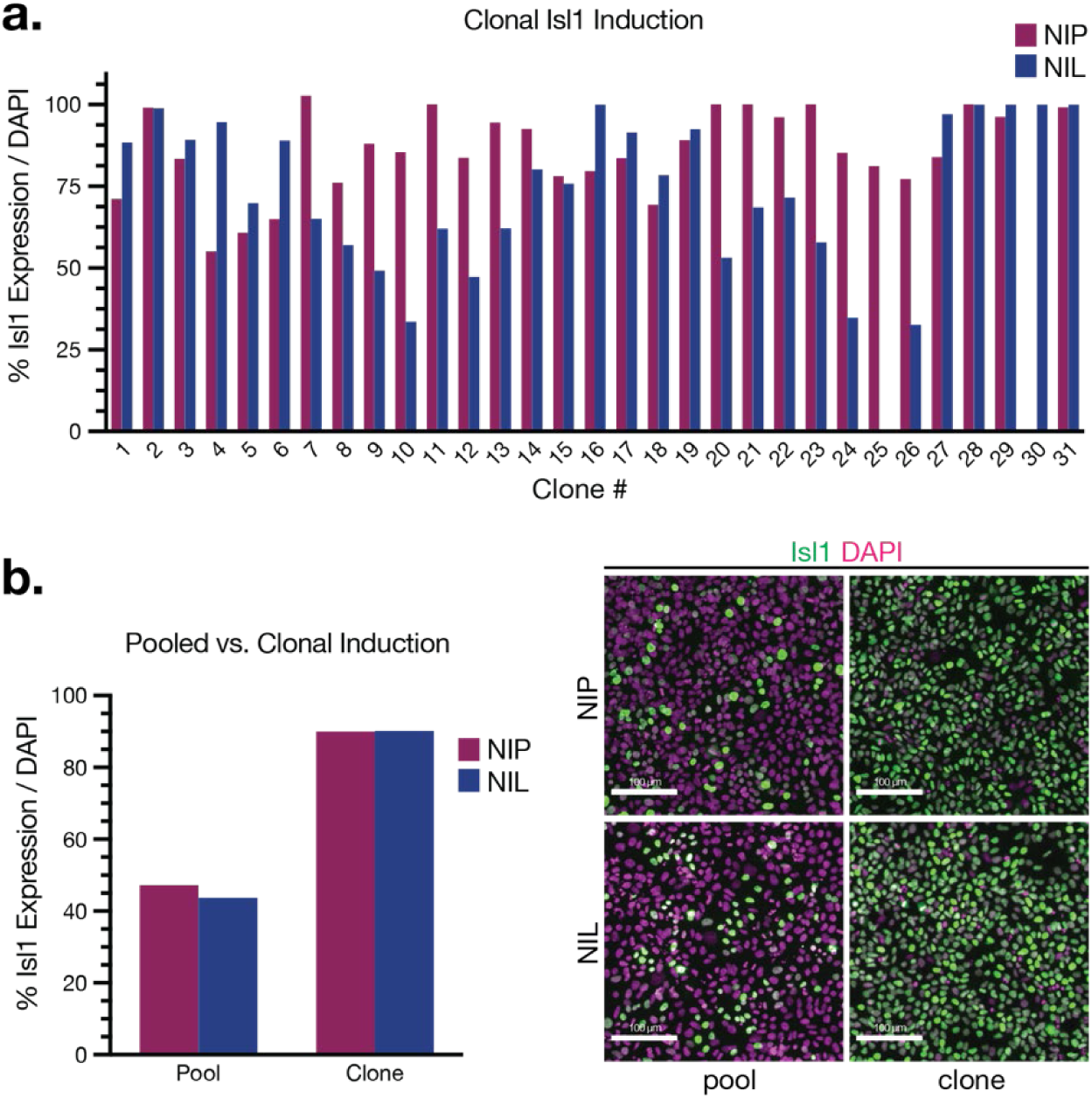
Clonal optimized induction of NIP and NIL iPSC programming. (**a**) Percent Islet1 (Isl1) expression over DAPI 2 days post doxycycline induction across 31 NIP (red) and NIL (blue) clones. (**b**) Optimized clonal induction rate is over 2-fold more efficient than pooled induction. Percent Isl1 expression over DAPI from NIP (red) and NIL (blue) pooled or top clones. Representative images (right) of Isl1 expression (green) and DAPI (blue) from a pool (left) or top clones (right) in NIP (top) and NIL (bottom). Scale bar = 100 µm.

**Supplemental Figure 2.**
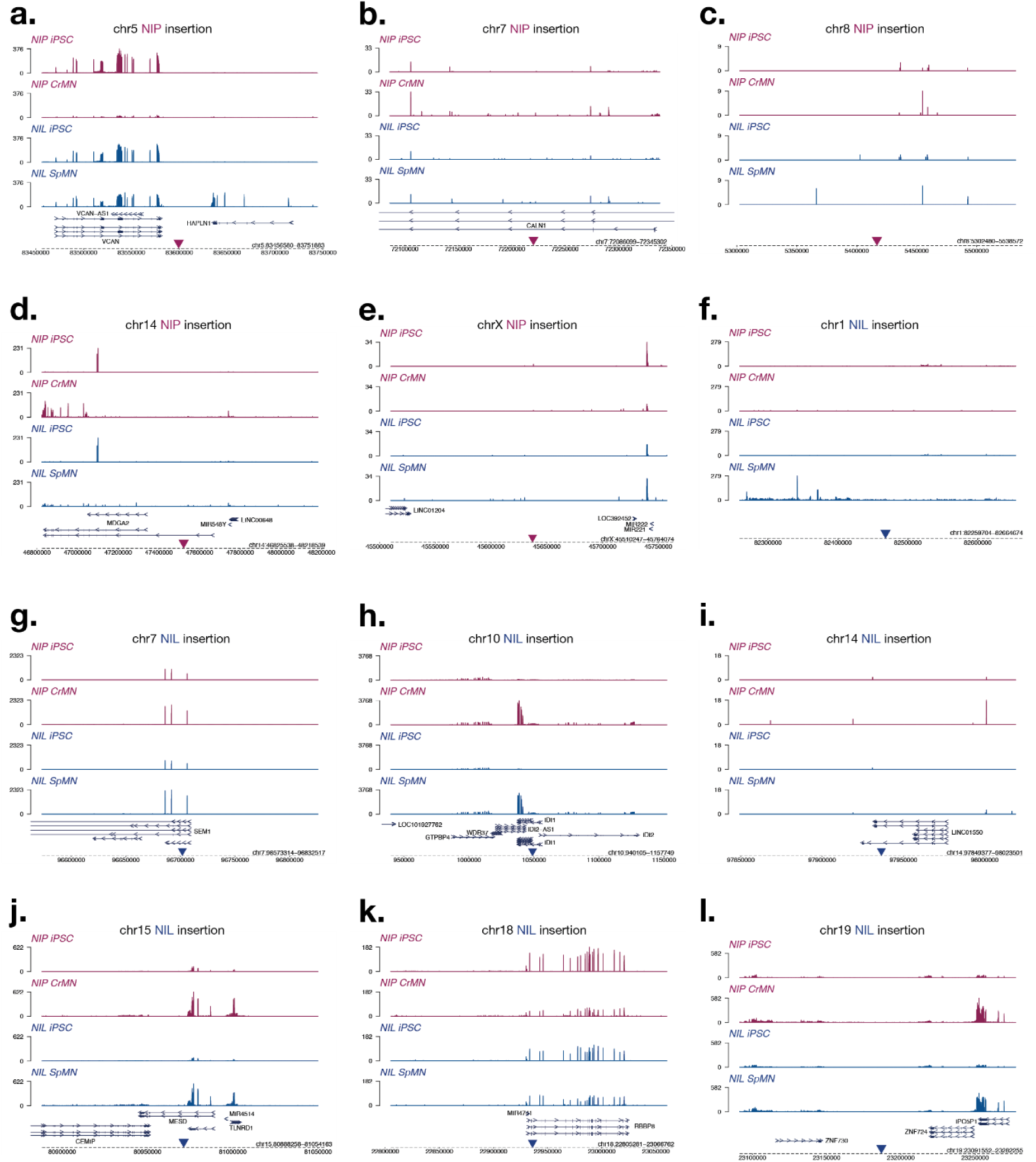
NIP and NIL genomic insertions have little effect on nearby transcription. (**a**-**l**) Normalized BigWig transcription near NIP cassette insertions in chromosomes (chr) 5 (**a**), 7 (**b**), 8 (**c**), 14 (**d**), and X (**e**) and NIL cassette insertions in chromosomes 1 (**f**), 7 (**g**), 10 (**h**), 14 (**i**), 15 (**j**), 18 (**k**), and 19 (**l**). Expression shown in NIP iPSCs (red, top), NIP CrMNs (red, second from top), NIL iPSCs (blue, second from bottom), and NIL SpMNs (blue, bottom). NIP insertion location = inverted red triangle. NIL insertion location = inverted blue triangle.

**Supplemental Figure 3.**
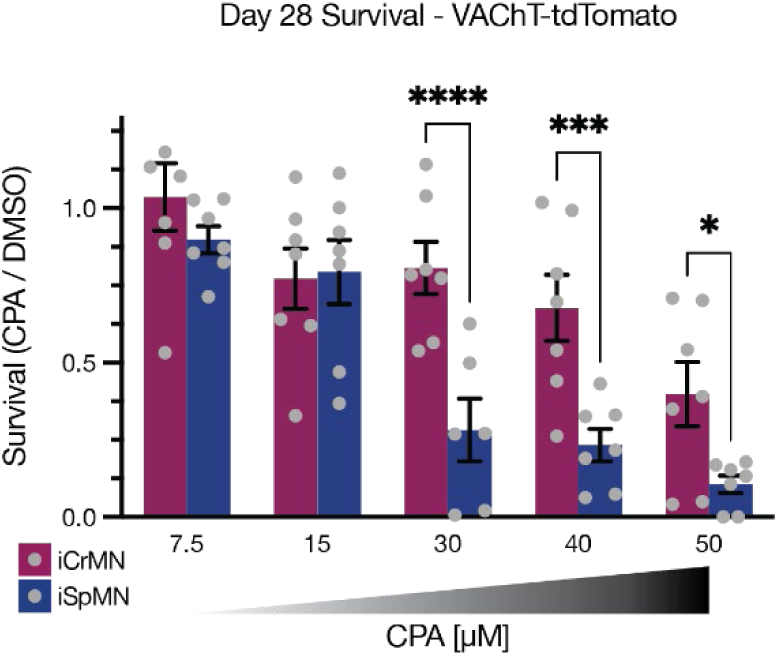
Measuring survival by imaging VAChT-tdT expressing recapitulates MTT cell viability. iCrMNs are more resistant than iSpMNs after 18 day CPA treatment. (n = 6-7, mean ± SEM, two-way ANOVA followed by Šídák’s multiple comparisons test). *p<0.0332, **p<0.0021, ***p<0.0002, ****p<0.0001.

**Supplemental Figure 4.**
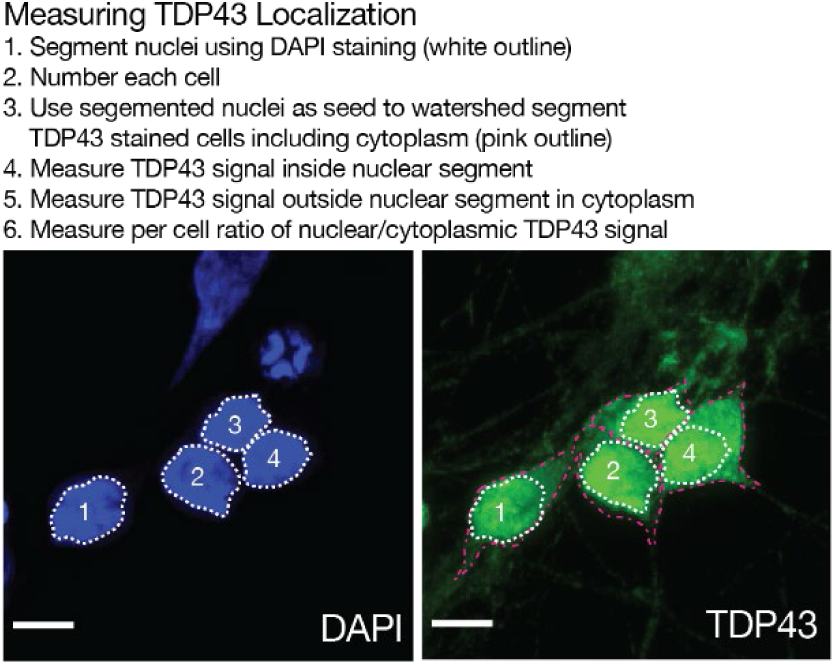
Method for measuring nuclear versus cytoplasmic localization of TDP43. Nuclei are segmented using DAPI (blue, right, white outlines) and individually numbered and used as seeds in the TDP43 channel (green, left) for watershed segmentation of cytoplasm per cell (magenta outlines). TDP43 fluorescent intensity in the nucleus is compared over intensity in the cytoplasm per cell.

**Supplemental Figure 5.**
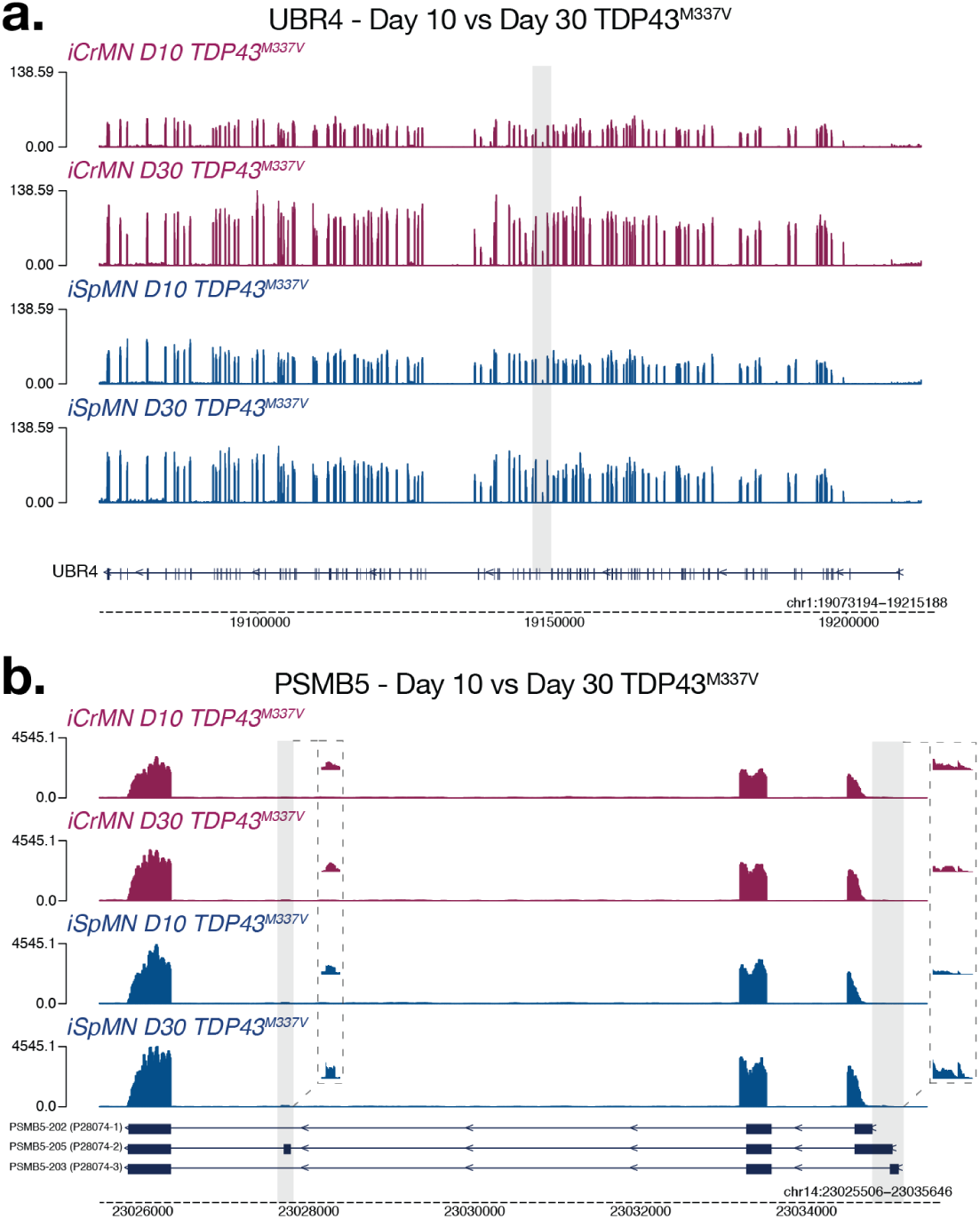
Differential splicing in TDP43^M337V^ iCrMNs and iSpMNs of UBR5 and PSMB5. (**a**-**b**) Normalized BigWig coverage plots for UBR5 (**a**) and PSMB5 (**b**). Expression shown in TDP43^M337V^ day 10 iCrMNs (red, top), day 30 iCrMNs (red, second from top), day 10 iSpMNs (blue, second from bottom), and day 30 iSpMNs (blue, bottom). Grey rectangles indicate locations of differential splicing identified by LeafCutter analysis.

**Supplemental Figure 6.**
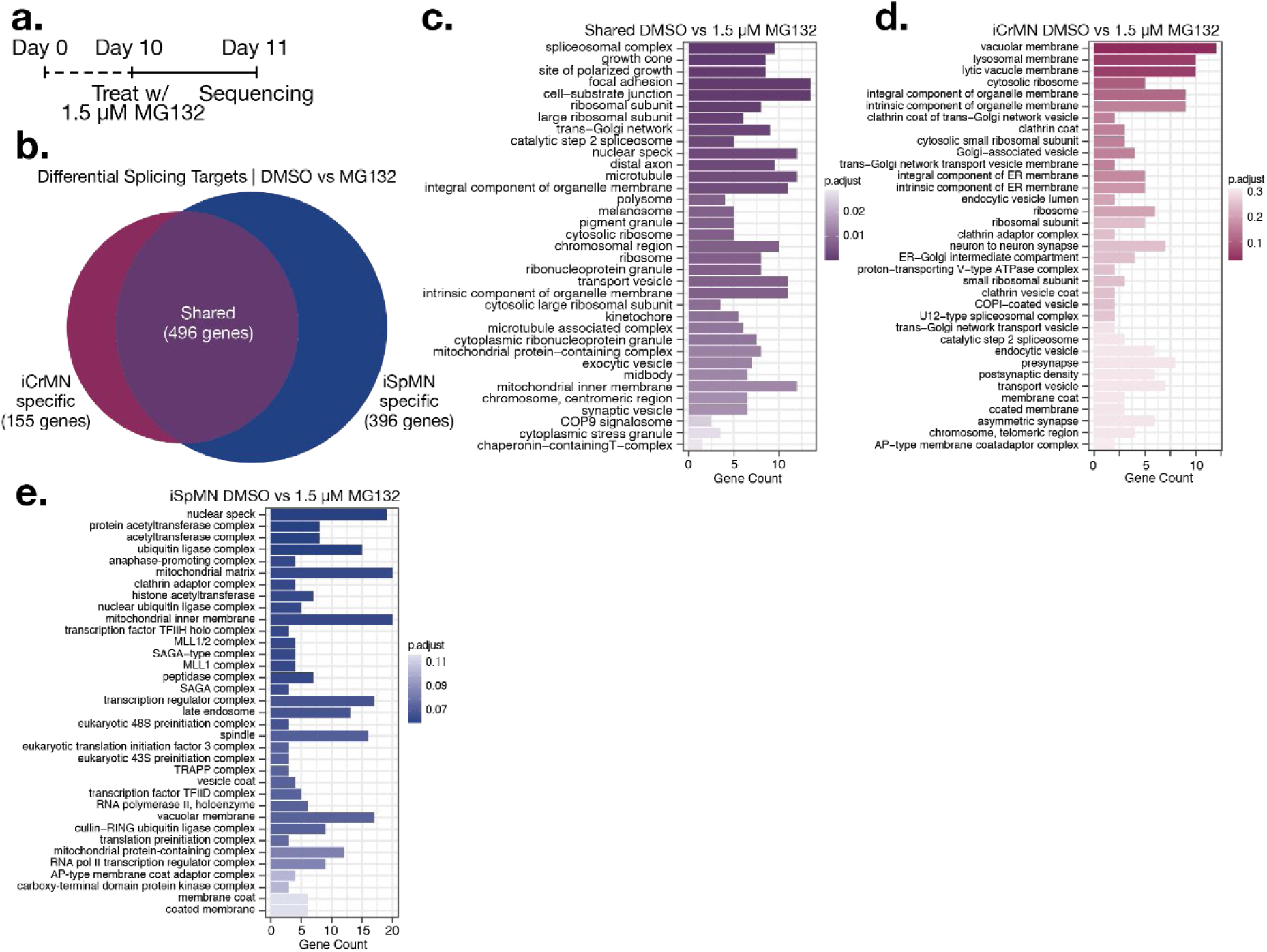
Cell type specific splicing changes after equivalent MG132 treatment in iCrMNs and iSpMNs. (**a**) Experimental outline: iCrMNs and iSpMNs are differentiated from day 0 to day 10 and treated with 1.5 MG132 on day 10 and collected 24 hours late for RNAseq. (**c**) DMSO versus MG132 DSGs specific to iCrMNs (155 genes), iSpMNs (396 genes), and shared (496 genes) after the removal of DSGs between DMSO treated iCrMNs and iSpMNs. (**d**, **e**, & **f**) Gene ontology (GO) enriched biological functions for shared (**d**) iCrMN-specific (**e**) and iSpMN-specific (**f**) DMSO vs MG132 DSGs.

**Supplemental Figure 7.**
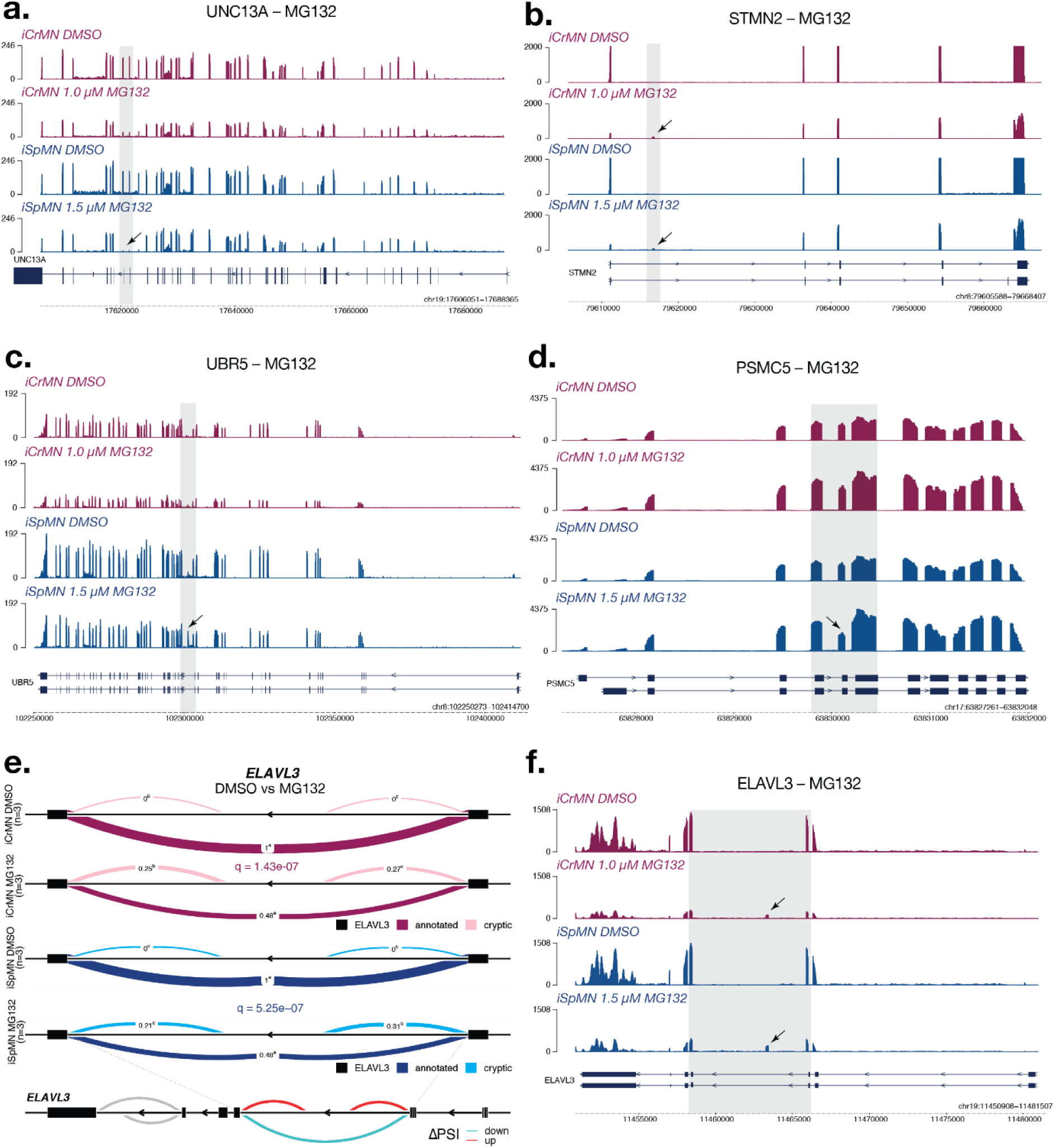
Synaptic, splicing, and proteostasis related splicing changes after MG132 induced TDP43 nuclear depletion in iCrMNs and iSpMNs. (**a**-**b**,**d**-**f**) Normalized BigWig coverage plots for UNC13A (**a**), STMN2 (**b**), ELAVL3 (**d**), UBR4 (**e**), and PSMC5 (**f**). Expression shown in DMSO treated iCrMNs (red, top), 1.0 µM MG132 treated iCrMNs (red, second from top), DMSO treated iSpMNs (blue, second from bottom), and 1.5 µM MG132 treated iSpMNs (blue, bottom). Grey rectangles indicate locations of differential splicing identified by LeafCutter analysis. Arrows indicate observable expression of differential splicing events. (**c**) LeafCutter analysis of differentially spliced intron clusters in the RNA-binding protein ELAVL3. Exons, black boxes. Red (iCrMN) and blue (iSpMN) band thickness and inserted values represent proportion of spliced exon-exon pairs. Change in percent spliced in (ΔPSI).

## Acknowledgements

We would like to thank current and former members of the Mazzoni lab for constructive comments and feedback, and in particular Milica Bulajić and Görkem Garipler for assistance with RNAseq library preparation. We thank Elizabeth Calder and Hynek Wichterle for advice and expertise for growing and differentiating human iPSCs to motor neuron fate. We thank Congyi Lu and Neville Sanjana for the iNeurog2/1 line and advice for differentiating iNs. We thank members of the Chan Zuckerberg Initiative Neurodegeneration Challenge Network for advice and expertise related to human iPSC lines and modeling ALS. We thank current and former staff in the NYU Biology Genomics Core in the Center for Genomics and Systems Biology for assistance and expertise in RNA sequencing and library preparation.

## Funding

DEI: NINDS F31NS103447

EOM: NIA R21AG067174, NYSDOH C32243GG, & CZI 2020-222014

## Competing Interest

The authors declare no competing interests.

## Author Contributions

Conceptualization: DEI, DA, EOM

Data curation: DEI, AT, EN, AB, SDE, TJ, SZ

Formal analysis: DEI, AT, EN, AB, SDE, TJ, SZ

Funding acquisition: DEI, EOM

Investigation: DEI, AT, EN, SDE, TJ, SZ

Methodology: DEI, AT, EN, AB, SDE, TJ, SZ, DA

Project administration: DEI, EOM

Resources: DEI, AB, SZ, DA, MW, EOM

Software: DEI, TJ, SZ

Supervision: DEI, EOM

Visualization: DEI, AT, EN, AB, SDE, TJ, SZ

Writing – original draft: DEI, EOM

Writing – review and editing: DEI, AT, EN, AB, SDE, TJ, SZ, DA, MW, EOM

## Notes

### Competing Interest Statement

The authors have declared no competing interest.

## References

1. Brown, R. H. & Al-Chalabi, A. Amyotrophic Lateral Sclerosis. http://dx.doi.org/10.1056/NEJMra1603471 377, 162–172 (2017).

2. Okamoto, K. et al. Oculomotor nuclear pathology in amyotrophic lateral sclerosis. Acta Neuropathologica 1993 85:5 85, 458–462 (1993).

3. Lawyer, T. & Netsky, M. G. AMYOTROPHIC LATERAL SCLEROSIS: A Clinicoanatomic Study of Fifty-Three Cases. A.M.A. Archives of Neurology & Psychiatry 69, 171–192 (1953).

4. Kanning, K. C., Kaplan, A. & Henderson, C. E. Motor Neuron Diversity in Development and Disease. Annu. Rev. Neurosci 33, 409–40 (2010).

5. Haenggeli, C. & Kato, A. C. Differential vulnerability of cranial motoneurons in mouse models with motor neuron degeneration. Neuroscience Letters 335, 39–43 (2002).

6. Allodi, I. et al. Modeling Motor Neuron Resilience in ALS Using Stem Cells. Stem Cell Reports 12, 1329–1341 (2019).

7. An, D. et al. Stem cell-derived cranial and spinal motor neurons reveal proteostatic differences between ALS resistant and sensitive motor neurons. eLife 8, (2019).

8. Yerbury, J. J., Farrawell, N. E. & Mcalary, L. Proteome Homeostasis Dysfunction: A Unifying Principle in ALS Pathogenesis. (2020) doi:10.1016/j.tins.2020.03.002.

9. Ruegsegger, C. & Saxena, S. Proteostasis impairment in ALS. Brain Research 1648, 571–579 (2016).

10. Kurtishi, A., Rosen, B., Patil, K. S., Alves, G. W. & Møller, S. G. Cellular Proteostasis in Neurodegeneration. Molecular Neurobiology 56, 3676–3689 (2019).

11. Hetz, C. & Mollereau, B. Disturbance of endoplasmic reticulum proteostasis in neurodegenerative diseases. Nature Reviews Neuroscience 2014 15:4 15, 233–249 (2014).

12. Bruijn, L. I. et al. Aggregation and Motor Neuron Toxicity of an ALS-Linked SOD1 Mutant Independent from Wild-Type SOD1. Science 281, 1851–1854 (1998).

13. Neumann, M. et al. Ubiquitinated TDP-43 in frontotemporal lobar degeneration and amyotrophic lateral sclerosis. Science 314, 130–133 (2006).

14. Deng, H.-X. et al. FUS-immunoreactive inclusions are a common feature in sporadic and non-SOD1 familial amyotrophic lateral sclerosis. Annals of Neurology 67, 739–748 (2010).

15. Zu, T. et al. RAN proteins and RNA foci from antisense transcripts in C9ORF72 ALS and frontotemporal dementia. Proceedings of the National Academy of Sciences 110, E4968–E4977 (2013).

16. Alami, N. H. et al. Axonal transport of TDP-43 mRNA granules in neurons is impaired by ALS-causing mutations. Neuron 81, 536 (2014).

17. Lee, E. B., Lee, V. M.-Y. & Trojanowski, J. Q. Gains or losses: molecular mechanisms of TDP43-mediated neurodegeneration. Nature Reviews Neuroscience 2011 13:1 13, 38–50 (2011).

18. Humphrey, J., Emmett, W., Fratta, P., Isaacs, A. M. & Plagnol, V. Quantitative analysis of cryptic splicing associated with TDP-43 depletion. BMC Medical Genomics 2017 10:1 10, 1–17 (2017).

19. Ling, J. P., Pletnikova, O., Troncoso, J. C. & Wong, P. C. TDP-43 repression of nonconserved cryptic exons is compromised in ALS-FTD. Science 349, 650–655 (2015).

20. Klim, J. R. et al. ALS-implicated protein TDP-43 sustains levels of STMN2, a mediator of motor neuron growth and repair. Nature Neuroscience 22, 167–179 (2019).

21. Melamed, Z. et al. Premature polyadenylation-mediated loss of stathmin-2 is a hallmark of TDP-43-dependent neurodegeneration. Nature Neuroscience 22, 180–190 (2019).

22. Brown, A.-L. et al. Common ALS/FTD risk variants in UNC13A exacerbate its cryptic splicing and loss upon TDP-43 mislocalization. bioRxiv 2021.04.02.438170 (2021) doi:10.1101/2021.04.02.438170.

23. Ma, X. R. et al. TDP-43 represses cryptic exon inclusion in FTD/ALS gene UNC13A. bioRxiv 2021.04.02.438213 (2021) doi:10.1101/2021.04.02.438213.

24. Amoroso, M. W. et al. Accelerated High-Yield Generation of Limb-Innervating Motor Neurons from Human Stem Cells. doi:10.1523/JNEUROSCI.0906-12.2013.

25. Maury, Y. et al. Combinatorial analysis of developmental cues efficiently converts human pluripotent stem cells into multiple neuronal subtypes. Nature Biotechnology 33, 89–96 (2015).

26. Aydin, B. & Mazzoni, E. O. Cell reprogramming: The many roads to success. Annual Review of Cell and Developmental Biology 35, 433–452 (2019).

27. Wang, H., Yang, Y., Liu, J. & Qian, L. Direct cell reprogramming: approaches, mechanisms and progress. Nature Reviews Molecular Cell Biology 2021 22:6 22, 410–424 (2021).

28. Mazzoni, E. O. et al. Synergistic binding of transcription factors to cell-specific enhancers programs motor neuron identity. Nature neuroscience 16, 1219–27 (2013).

29. Iacovino, M. et al. Inducible cassette exchange: A rapid and efficient system enabling conditional gene expression in embryonic stem and primary cells. Stem Cells 29, 1580–1587 (2011).

30. De Santis, R. et al. Direct conversion of human pluripotent stem cells into cranial motor neurons using a piggyBac vector. Stem Cell Research 29, 189–196 (2018).

31. Lu, C. et al. Overexpression of NEUROG2 and NEUROG1 in human embryonic stem cells produces a network of excitatory and inhibitory neurons. FASEB Journal 33, 5287–5299 (2019).

32. Garcia-Diaz, A. et al. Standardized Reporter Systems for Purification and Imaging of Human Pluripotent Stem Cell-derived Motor Neurons and Other Cholinergic Cells. Neuroscience 450, 48–56 (2020).

33. Maury, Y. et al. Combinatorial analysis of developmental cues efficiently converts human pluripotent stem cells into multiple neuronal subtypes. Nature Biotechnology 33, 89–96 (2015).

34. Love, M. I., Huber, W. & Anders, S. Moderated estimation of fold change and dispersion for RNA-seq data with DESeq2. Genome Biology 2014 15:12 15, 1–21 (2014).

35. Jessell, T. M. Neuronal specification in the spinal cord: inductive signals and transcriptional codes. Nature Reviews Genetics 1, 20–9 (2000).

36. Song, M.-R. et al. T-Box transcription factor Tbx20 regulates a genetic program for cranial motor neuron cell body migration. Development (Cambridge, England) 133, 4945–4955 (2006).

37. Müller, M., Jabs, N., Lorke, D. E., Fritzsch, B. & Sander, M. Nkx6.1 controls migration and axon pathfinding of cranial branchio-motoneurons. Development 130, 5815–5826 (2003).

38. Prakash, N. et al. Nkx6-1 controls the identity and fate of red nucleus and oculomotor neurons in the mouse midbrain. Development (Cambridge, England) 136, 2545 (2009).

39. Bjorke, B. et al. Contralateral migration of oculomotor neurons is regulated by Slit/Robo signaling. Neural Development 11, (2016).

40. Partanen, J. FGF signalling pathways in development of the midbrain and anterior hindbrain. Journal of Neurochemistry 101, 1185–1193 (2007).

41. Yaylaoglu, M. B. et al. Comprehensive expression atlas of fibroblast growth factors and their receptors generated by a novel robotic in situ hybridization platform. Developmental Dynamics 234, 371–386 (2005).

42. Hattori, Y., Yamasaki, M., Konishi, M. & Itoh, N. Spatially restricted expression of fibroblast growth factor-10 mRNA in the rat brain. Molecular Brain Research 47, 139–146 (1997).

43. Grillet, N., Dubreuil, V., Dufour, H. D. & Brunet, J. F. Dynamic Expression of RGS4 in the Developing Nervous System and Regulation by the Neural Type-Specific Transcription Factor Phox2b. Journal of Neuroscience 23, 10613–10621 (2003).

44. Magen, I. et al. Circulating miR-181 is a prognostic biomarker for amyotrophic lateral sclerosis. Nature Neuroscience 2021 24:11 24, 1534–1541 (2021).

45. Matus, S., Valenzuela, V., Medinas, D. B. & Hetz, C. ER dysfunction and protein folding stress in ALS. International Journal of Cell Biology (2013) doi:10.1155/2013/674751.

46. Merlie, J. P., Sebbane, R., Tzartos, S. & Lindstrom, J. Inhibition of glycosylation with tunicamycin blocks assembly of newly synthesized acetylcholine receptor subunits in muscle cells. Journal of Biological Chemistry 257, 2694–2701 (1982).

47. Uyama, Y., Imaizumi, Y. & Watanabe, M. Cyclopiazonic acid, an inhibitor of Ca2+-ATPase in sarcoplasmic reticulum, increases excitability in ileal smooth muscle. British Journal of Pharmacology 110, 565–572 (1993).

48. Thams, S. et al. A Stem Cell-Based Screening Platform Identifies Compounds that Desensitize Motor Neurons to Endoplasmic Reticulum Stress. Molecular Therapy 27, 87–101 (2019).

49. Mosmann, T. Rapid colorimetric assay for cellular growth and survival: Application to proliferation and cytotoxicity assays. Journal of Immunological Methods 65, 55–63 (1983).

50. Dantuma, N. P., Bott, L. C. & Mcallister, A. K. The ubiquitin-proteasome system in neurodegenerative diseases: precipitating factor, yet part of the solution. (2014) doi:10.3389/fnmol.2014.00070.

51. Carpenter, A. E. et al. CellProfiler: image analysis software for identifying and quantifying cell phenotypes. Genome biology 7, R100 (2006).

52. Kisselev, A. F. & Goldberg, A. L. Proteasome inhibitors: from research tools to drug candidates. Chemistry & Biology 8, 739–758 (2001).

53. Goldberg, A. L. Development of proteasome inhibitors as research tools and cancer drugs. 199, (2012).

54. Ruggeri, B., Miknyoczki, S., Dorsey, B. & Hui, A. M. The Development and Pharmacology of Proteasome Inhibitors for the Management and Treatment of Cancer. Advances in pharmacology (San Diego, Calif.) 57, 91–135 (2009).

55. Sin, N. et al. Total synthesis of the-potent proteasome inhibitor epoxomicin: a useful tool for understanding proteasome biology. Bioorganic & Medicinal Chemistry Letters 9, 2283–2288 (1999).

56. Ling, S. C., Polymenidou, M. & Cleveland, D. W. Converging Mechanisms in ALS and FTD: Disrupted RNA and Protein Homeostasis. Neuron 79, 416–438 (2013).

57. Li, Y. I. et al. Annotation-free quantification of RNA splicing using LeafCutter. Nature Genetics 2017 50:1 50, 151–158 (2017).

58. Choi, K. D. et al. Genetic Variants Associated with Episodic Ataxia in Korea. Scientific Reports 2017 7:1 7, 1–11 (2017).

59. Conroy, J. et al. A novel locus for episodic ataxia:UBR4 the likely candidate. European Journal of Human Genetics 2014 22:4 22, 505–510 (2013).

60. Bieler, S. et al. Low Dose Proteasome Inhibition Affects Alternative Splicing. (2012) doi:10.1021/pr300435c.

61. Huang, H. H. et al. Proteasome inhibitor-induced modulation reveals the spliceosome as a specific therapeutic vulnerability in multiple myeloma. Nature Communications 2020 11:1 11, 1–14 (2020).

62. Koyuncu, S. et al. The ubiquitin ligase UBR5 suppresses proteostasis collapse in pluripotent stem cells from Huntington’s disease patients. Nature Communications 2018 9:1 9, 1–22 (2018).

63. Wang, Q., Conlon, E. G., Manley, J. L. & Rio, D. C. Widespread intron retention impairs protein homeostasis in C9orf72 ALS brains. Genome Research 31, 1705–1715 (2020).

64. Diaz-Garcia, S. et al. Nuclear depletion of RNA-binding protein ELAVL3 (HuC) in sporadic and familial amyotrophic lateral sclerosis. Acta Neuropathologica 2021 1, 1–17 (2021).

65. Ichida, J. K. & Ko, C. P. Organoids Develop Motor Skills: 3D Human Neuromuscular Junctions. Cell Stem Cell 26, 131–133 (2020).

66. Faustino Martins, J. M., et al. Self-Organizing 3D Human Trunk Neuromuscular Organoids. Cell Stem Cell 26, 172–186.e6 (2020).

67. Andersen, J. et al. Generation of Functional Human 3D Cortico-Motor Assembloids. Cell 183, 1913–1929.e26 (2020).

68. Pediaditakis, I. et al. Modeling alpha-synuclein pathology in a human brain-chip to assess blood-brain barrier disruption. Nature Communications 2021 12:1 12, 1–17 (2021).

69. Hetz, C. et al. XBP-1 deficiency in the nervous system protects against amyotrophic lateral sclerosis by increasing autophagy. Genes and Development 23, 2294–2306 (2009).

70. Atkin, J. D. et al. Endoplasmic reticulum stress and induction of the unfolded protein response in human sporadic amyotrophic lateral sclerosis. Neurobiology of Disease 30, 400–407 (2008).

71. Lee, S. et al. Activation of HIPK2 Promotes ER Stress-Mediated Neurodegeneration in Amyotrophic Lateral Sclerosis. Neuron 41–55 (2016) doi:10.1016/j.neuron.2016.05.021.

72. Saxena, S., Cabuy, E. & Caroni, P. A role for motoneuron subtype-selective ER stress in disease manifestations of FALS mice. Nature neuroscience 12, 627–636 (2009).

73. Kaplan, A. et al. Neuronal Matrix Metalloproteinase-9 Is a Determinant of Selective Neurodegeneration. Neuron 81, 333–348 (2014).

74. Allodi, I. et al. Differential neuronal vulnerability identifies IGF-2 as a protective factor in ALS. Scientific Reports 2016 6:1 6, 1–14 (2016).

75. Lee, H. et al. Multi-omic analysis of selectively vulnerable motor neuron subtypes implicates altered lipid metabolism in ALS. Nature Neuroscience 2021 1–13 (2021) doi:10.1038/s41593-021-00944-z.

76. Arnold, E. S. et al. ALS-linked TDP-43 mutations produce aberrant RNA splicing and adult-onset motor neuron disease without aggregation or loss of nuclear TDP-43. Proceedings of the National Academy of Sciences of the United States of America 110, (2013).

77. Jone Ló pez-Erauskin, A., et al. ALS/FTD-Linked Mutation in FUS Suppresses Intra-axonal Protein Synthesis and Drives Disease Without Nuclear Loss-of-Function of FUS. Neuron 100, 816–830.e7 (2018).

78. Fratta, P. et al. Mice with endogenous TDP-43 mutations exhibit gain of splicing function and characteristics of amyotrophic lateral sclerosis. The EMBO Journal 37, e98684 (2018).

79. Humphrey, J. et al. FUS ALS-causative mutations impair FUS autoregulation and splicing factor networks through intron retention. Nucleic Acids Research 48, 6889–6905 (2020).

80. Butti, Z. & Patten, S. A. RNA dysregulation in amyotrophic lateral sclerosis. Frontiers in Genetics 10, 712 (2019).

81. Perrone, B. et al. Alternative Splicing of ALS Genes: Misregulation and Potential Therapies. Cellular and Molecular Neurobiology 2019 40:1 40, 1–14 (2019).

82. Deshaies, J. E. et al. TDP-43 regulates the alternative splicing of hnRNP A1 to yield an aggregation-prone variant in amyotrophic lateral sclerosis. Brain 141, 1320–1333 (2018).

83. Torres, P. et al. Cryptic exon splicing function of TARDBP interacts with autophagy in nervous tissue. Autophagy 14, 1398–1403 (2018).

84. Pantazis, C. B. et al. A reference induced pluripotent stem cell line for large-scale collaborative studies. bioRxiv 20, 2021.12.15.472643 (2021).

85. Garone, M. G. et al. Conversion of Human Induced Pluripotent Stem Cells (iPSCs) into Functional Spinal and Cranial Motor Neurons Using PiggyBac Vectors Conversion of Human Induced Pluripotent Stem Cells (iPSCs) into Functional Spinal and Cranial Motor Neurons Using PiggyBac. J. Vis. Exp 59321 (2019) doi:10.3791/59321.

86. Dobin, A. & Gingeras, T. R. Optimizing RNA-Seq Mapping with STAR. Methods in Molecular Biology 1415, 245–262 (2016).

87. Li, H. et al. The Sequence Alignment/Map format and SAMtools. Bioinformatics 25, 2078–2079 (2009).

88. Danecek, P. et al. Twelve years of SAMtools and BCFtools. GigaScience 10, 1–4 (2021).

89. Ramírez, F., Dündar, F., Diehl, S., Grüning, B. A. & Manke, T. deepTools: a flexible platform for exploring deep-sequencing data. Nucleic Acids Research 42, W187 (2014).

90. Liao, Y., Smyth, G. K. & Shi, W. FeatureCounts: An efficient general purpose program for assigning sequence reads to genomic features. Bioinformatics 30, 923–930 (2014).

91. Liao, Y., Smyth, G. K. & Shi, W. The R package Rsubread is easier, faster, cheaper and better for alignment and quantification of RNA sequencing reads. Nucleic Acids Research 47, e47–e47 (2019).

92. Pohl, A. & Beato, M. bwtool: a tool for bigWig files. Bioinformatics 30, 1618–1619 (2014).

93. Yu, G., Wang, L. G., Yan, G. R. & He, Q. Y. DOSE: an R/Bioconductor package for disease ontology semantic and enrichment analysis. Bioinformatics 31, 608–609 (2015).

94. Yu, G. enrichplot: Visualization of Functional Enrichment Result. R package version 1.14.1 (2021).

